# Learning-induced plasticity decreases cortical engram cell dendritic excitability during memory recall

**DOI:** 10.1101/2025.01.08.631892

**Authors:** Marius Rosier, George Stuyt, Luca Godenzini, Tomás J. Ryan, Lucy M. Palmer

## Abstract

The ability to associate stimuli to create a memory is one of the most fundamental functions of the brain. Research from the past decade has revealed that memory is encoded in sparse neuronal networks active during learning called engram cells. Although the cortex is recognized as playing an important role in memory, the biophysical properties of cortical engram cells are largely unknown. To address this, we tagged engram cells in the auditory cortex during tone fear conditioning and compared their dendritic and somatic properties with neighbouring non-engram cells. Using two-photon calcium imaging, we show that tuft dendrites of engram cells had dampened, but synchronous, activity during recall. *Ex vivo* patch-clamp recordings illustrated that engram cells were preferentially connected with neighbouring engram cells and had decreased excitability due to a transient increase in Ih current. Together, these findings reveal Ih-driven intrinsic plasticity which leads to specific information processing in engram cells.

## Introduction

Memories are stored as ensembles of synaptically-connected neurons called engram cells^1,2^, that can be identified using molecular tagging based on immediate early gene or calcium activity^3–6^. The allocation of engram cells to the memory trace is believed to be determined by their activity at the time of learning^7,8^. Engram cells then undergo intrinsic and synaptic plasticity mechanisms during consolidation^1,9,10^ that play an important role during memory recall^11–15^.

Previous research has identified populations of engram cells in several cortical regions^12,14,16–18^. The cortex has a hierarchical structure which is well suited for the association of distinct information pathways required for memory formation and recall, receiving both feedback (internal) and feedforward (external) information^19–22^. Of particular importance for associative learning such as fear conditioning^23–26^, the upper layers of the cortex are the target of long-range feedback input^22^. Residing in cortical layer 1, the tuft dendrites of pyramidal neurons are therefore an ideal candidate for influencing memory processes and have been shown to have increased activity following tone fear conditioning^27^. By transforming synaptic input into large non-linear dendritic voltage events called dendritic spikes, dendrites can directly and potently influence somatic output^28,29^. Therefore, the modulation of dendritic excitability is an effective and rapid mechanism to dynamically adapt to incoming patterns of synaptic activity. However, whether learning-induced intrinsic plasticity occurs in engram cells at the dendritic level remains to be investigated. Of particular importance to the excitability of distal tuft dendrites are the hyperpolarization-activated cyclic nucleotide-gated (HCN) channels which generate an inward voltage-gated current (Ih)^30^. Known to decrease dendritic excitability^31,32^, Ih regulates synaptic plasticity^33,34^ and can directly influence the generation of dendritic spikes^35,36^. Furthermore, Ih is critical for the synchronization of neuronal activity^36–38^ which is an important driver of brain function and communication^22^. Prone to plasticity following learning^39,40^, Ih has been shown to play an important role in memory processes^41,42^ and may therefore play a defining role in the function of engram cells by directly and rapidly modulating dendritic excitability.

Here, we identify engram cells forming a network within the deep layers of the auditory cortex. Using *in vivo* calcium imaging, we reveal a decreased, but synchronized, pattern of activity in the tuft dendrites of cortical engram cells during memory recall. Consistent with this, *ex vivo* patch clamp recordings illustrate that engram cells have decreased excitability compared to neighbouring non-engram cells, which was due to increased Ih currents. These learning-induced changes in cortical engram cells were transient, as cellular excitability had returned to pre-conditioned levels after 1 week. Our findings highlight the flexible and defined features of engram cells in the cortex and shed light on the sensory cortex as playing an important role in memory processes.

## Results

### Engram cells in the deep layers of the auditory cortex

To label engram cells, we injected AAV_9-_TRE-mCherry into the auditory cortex of transgenic mice expressing the tetracycline transactivator (tTA) under the control of the immediate early gene *c-fos* promoter (c-fos-tTA; Figure 1a)^3,43^. Binding of the tTA to the TRE promoter drives the expression of the mCherry transgene in cells with c-Fos expression. This labelling of engram cells is under the control of doxycycline (DOX), and mice are kept on a DOX-containing diet (40 mg/kg) to prevent labelling prior to learning (Figure 1a). 48 h prior to engram labelling, mice were taken off the DOX-containing diet (OFF DOX) to enable the expression of the mCherry transgene. Mice were then fear conditioned by 5 pairings of an ascending tone (5-15 KHz, 30 s, 70 dB)^24^ which co-terminated with an electric foot shock (2 sec, 0.6 mA; CS+). Immediately following fear conditioning, mice were placed back ON DOX to cease labelling. To reduce nonspecific labelling, mice were placed in a quiet environment during the OFF DOX period and for 24 h following fear conditioning (Figure S1). First, to assess learning-induced labelling of putative engram cells, we compared the number of mCherry-positive neurons between fear conditioned and home cage animals, which remained in their home cage during the OFF DOX period (Figure 1b). Animals were perfused 24 h following the end of the OFF DOX period, and the number of mCherry neurons was quantified as a percentage of the total number of cells (according to DAPI staining). In contrast to layer 2/3 pyramidal neurons, the number of neurons labelled in layers 5 and 6 was significantly greater in fear conditioned mice (3.39 ± 0.32 %; n = 9) compared to home cage mice (2.18 ± 0.41 % mCherry in DAPI; n = 5; Figure 1b-c and S2; p = 0.042). These findings illustrate learning-induced labelling of neurons in the deep layers of the auditory cortex. Since reactivation during recall is a key characteristic of engram cells^2^, we next assessed the recall-induced activity of neurons that were labelled during fear learning by evaluating c-Fos expression in mCherry-labelled neurons after recall. To achieve this, mice were exposed to the conditioning tone 24 h after fear conditioning and perfused 1 h later to quantify recall-induced expression of c-Fos (Figure 1d-e). Exposure to the CS+ elicited significant freezing behaviour during tone exposure and the inter-stimuli intervals (Figure 1d). Following recall, there was significantly greater c-Fos expression (24.95 ± 2.07 % c-Fos in DAPI; n = 6) and c-Fos/mCherry overlap (79.8 ± 4.2 % c-Fos in mCherry; n = 4) in neurons within layers 5 and 6 of the auditory cortex compared to neurons from mice that were not exposed to CS+ recall (12.95 ± 1.65 % c-Fos in DAPI, p = 0.001; 51.3 ± 8.1 % c-Fos in mCherry; n = 5; p = 0.032; Figure 1e). These results illustrate a recall-induced activation of neurons that were active during learning in the deep layers of the auditory cortex. Recall-induced activation of the auditory cortex was also observed in the superficial layers, however there was no significant c-Fos/mCherry overlap (77.41 ± 7.02 % (Recall, n = 5) vs 53.67 ± 8.09 (No Recall, n = 5); p = 0.11; Figure S1), suggesting these neurons were not selectively active during recall. Similar results were also reported when engram cells were labelled with EYFP-ChR2 (AAV_9-_TRE-EYFP-ChR2; Figure S1). Combined, these findings suggest a specific recall-induced reactivation of layer 5 and 6 neurons within the auditory cortex, which supports the classification of these labelled deep layer pyramidal neurons as engram cells.

**Figure 1.**
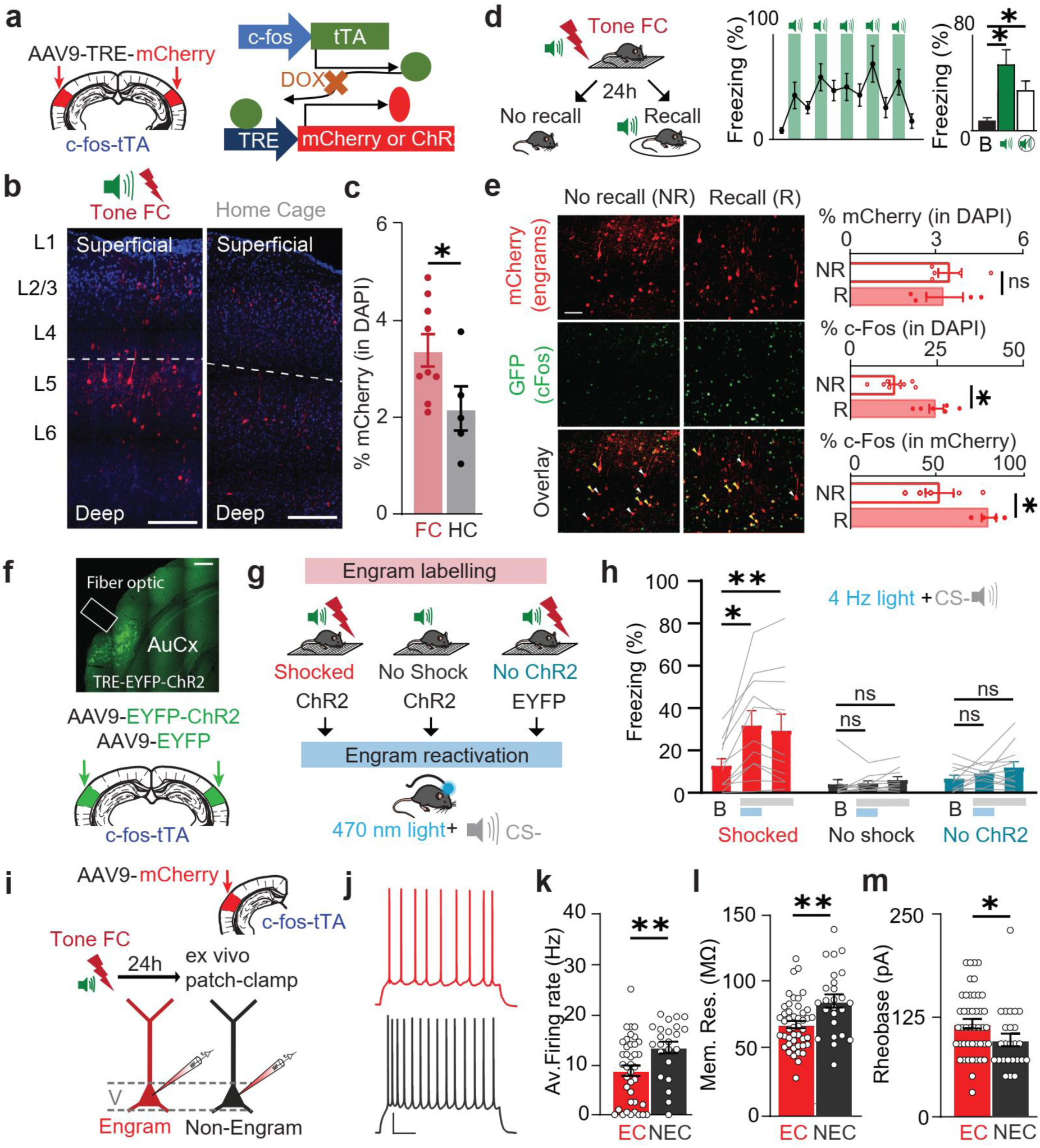
Engram cells populations in the deep layers of the auditory cortex with reduced intrinsic excitability. **a,** Engram labelling strategy using the c-fos-tTA system. An AAV with the TRE promoter was injected in the auditory cortex of fos-tTA transgenic mice to express a transgene under the c- fos promoter. **b,** Example pictures of engram labelling in the auditory cortex in a fear conditioned animal (left) and a home cage animal (right). Scale bars represent 100 µm. **c,** Quantification of mCherry cells on DAPI, revealing an increased number of cells labelled within the deep layers of the auditory cortex in conditioned animals (Tone FC; n = 9) compared to home cage (HC; n = 4). **d,** Fear recall was elicited by CS+ exposure 24h after conditioning (Recall), inducing freezing behaviour during tones (green bars) and the inter-stimuli interval (white bars). Animals were perfused 1h after recall. Animals from the No recall group remained in their home cage between conditioning and perfusion. **e,** Left: example pictures of mCherry (top), c-Fos (middle) and their overlay (bottom) for the Recall and No Recall groups. White arrows indicate mCherry+ neurons (non-reactivated) and yellow arrows the mCherry+/c-Fos+ neurons (reactivated). Scale bar represents 50 µm. Right: quantification of mCherry-expressing (top), c-Fos-expressing (middle) and mCherry/c-Fos expressing neurons. Tone recall 24h after conditioning and significantly induced engram cells reactivation in the deep layers as shown by the c-Fos/mCherry overlap increase in the Recall group (n = 4) vs no recall (n = 5). **f,** Representative histological section of the auditory cortex showing engram labelling with the AAV9-TRE-EYFP-ChR2 construct and the fiber optic cannula placement. Scale bar represents 500 µm. **g,** Experimental design and groups for the gain of function experiment. Light delivery was combined with a non-conditioned stimulus (CS-) during optogenetic sessions. **h,** Bar graph showing the average freezing quantification over 2 days of the same condition (4Hz combined with CS-) for all groups. Blue bars under the graph represent light ON, and grey bars the CS-. Significant freezing was elicited in the Shock group (n = 11) when light delivery was combined with the CS-. No freezing was observed in the No Shock group (n = 10) or the No ChR2 group (n = 12). **i,** Schematic of the experimental strategy. Auditory cortex slices were processed 24h following conditioning for patch-clamp recordings in slices. **j,** Example traces of action potential firing in an engram cell (EC; top, red) and a non-engram cell (NEC; bottom, grey) in response to a 150 pA current step injection. Scale bars represent 20mV and 200 ms. **k,** Average firing rate of EC (n = 46 from 30 mice) and NEC (n = 25 from 22 mice) between 130-150 pA current step injections, showing decreased firing rate in EC. **l,** Membrane resistance of EC and NEC, showing a decreased membrane resistance in EC. **m,** Rheobase of EC and NEC, showing an increased rheobase in EC. Data are represented as mean ± SEM and significance was assessed with two-tailed unpaired Mann-Whitney tests for e (*: p < 0.05), two-way ANOVA with a Dunnett correction for h (*: p < 0.05, **: p < 0.01), and two-tailed unpaired t-tests for k, l and m (*: p < 0.05, **: p < 0.01).

We next tested whether the reactivation of engram cells could elicit learned fear behaviour (freezing). The auditory cortex of c-fos-tTA transgenic mice were injected with either 1) AAV_9-_ TRE-ChR2-EYFP and conditioned with CS+ (Shocked, n = 11), 2) AAV_9-_TRE-ChR2-EYFP and exposed to the CS+ tone without pairing with an electric shock (No shock, n = 10), or 3) AAV_9-_TRE-EYFP (without the ChR2 transgene) and conditioned with CS+ (No ChR2, n = 12; Figure 1g). First, fear memory was assessed 24 h after conditioning by exposing animals to the CS+ (ascending tone; 5-15 KHz, 30 s, 70 dB) as well as a non-conditioned stimulus (CS-; white noise; 5-15 KHz, 30 s, 70 dB). Following fear conditioning, mice had higher rates of freezing to the CS+ compared to the CS-, suggesting that they were able to discriminate between the two stimuli (Figure S2c). Next, we tested whether photoactivation of engram cells can induce behavioural expression of fear memory (freezing) by delivering blue light (470 nm; 20 Hz or 4 Hz for 6 epochs of 1 mn) to the auditory cortex (Figure 1f and S4a). To test for the importance of salient sensory inputs, we combined the optogenetic reactivation with the CS- during some sessions (Figure S2). Here, the combination of 4 Hz photo-activation of engram cells and CS- exposure caused a significant increase in the freezing rate in the shocked mice (12.6 ± 3.5 % (Baseline) vs 31.5 ± 7.2 % (LED ON + CS), p = 0.004; Figure 1g). In contrast, there was no effect on freezing in control mice which either did not receive any shock (No shock: 3.8 ± 2.4 % (Baseline) vs 4.3 ± 1.3 (LED ON + CS), p = 0.99) or were not transfected with an opsin (No ChR2: 6.6 ± 1.7 % (Baseline) vs 8.9 ± 1.4 (LED ON + CS), p = 0.98; Figure 1h; Figure S2). Similar to the freezing during natural recall staying elevated during the inter-stimulus interval (Figure 1d), engram photoactivation (LED ON) in Shocked mice resulted in behavioural freezing that was long lasting and continued in the absence of LED (29.2 ± 8 % (LED OFF + CS), p = 0.01 vs Baseline; Figures 1h and S5, Table S1). Results for each session of optogenetic reactivation are shown in Figure S2, and the corresponding p-values are shown in Table S1. Taken together, cells labelled in the auditory cortex are defined as engram cells as they 1) are activated during learning and recall, and 2) can induce behavioural expression of memory when reactivated.

To investigate the properties of engram and non-engram cells, c-fos-tTA mice were injected with an AAV_9_-TRE-mCherry in the auditory cortex and engram cells were labelled during tone fear conditioning as described previously. 24 h after conditioning, *ex vivo* patch-clamp recordings were performed from layer 5 (L5) pyramidal engram (n = 46 from 30 mice) and non-engram (n = 25 from 22 mice) cells (Figure 1i). Similar to engram cells within the hippocampus^11^, cortical engram cells have increased spontaneous synaptic inputs frequency and amplitude compared to non-engram cells (Figure S3). Furthermore, compared to non-engram cells, engram cells had decreased intrinsic excitability during low input, with a significant decrease in firing rate (8.76 ± 1.05 vs 13.39 ± 1.17 Hz; p = 0.007; Figure 1j-k and S6), membrane resistance (68.71 ± 2.77 vs 86.44 ± 4.85 MΩ; p = 0.001; Figure 1l), and an increased rheobase (114.90 ± 5.90 vs 94.80 ± 7.77 pA; p = 0.044; Figure 1m). However, in response to high input patterns, engram cells had a significantly increased firing rate and burst firing (Figure S3), illustrating their dynamic responses to different input patterns. These findings illustrate engram cells have different intrinsic excitability compared to neighbouring non-engram cells, suggesting they may differentially encode behaviourally relevant information.

### Engram dendrites have decreased responses to conditioned stimuli during memory recall

Since the upper cortical layers, where the tuft dendrites of pyramidal neurons reside, are crucial for memory processes^23–25^, we assessed the activity of the tuft dendrites from engram and non-engram cells within the auditory cortex following fear learning. To achieve this, both AAV_9_-TRE-mCherry and AAV_1_-syn-jGCaMP7f were injected into the auditory cortex, and engram cells were labelled during auditory fear conditioning as described previously (Figure 2a). 24 h following fear conditioning, two-photon calcium activity was performed from the tuft dendrites of L5 pyramidal neurons during interleaved exposure to CS+ and CS- (Figure 2a). In response to the conditioned stimulus, there was a significant decrease in the rate of calcium events evoked in engram cell dendrites (baseline 0.06 ± 0.02 Hz vs CS+ 0.03 ± 0.01; p = 0.005; n = 27 from 7 mice; Figure 2c, e). In contrast, there was a significant increase in the dendritic calcium activity in response to CS+ in non-engram cells (baseline 0.04 ± 0.01 Hz vs CS+ 0.07 ± 0.01; p = 0.0349; n = 106 from 7 mice; Figure 2d, f). Despite the change in the evoked rate, there was no significant difference in the amplitude of the evoked calcium events in either engram cell or non-engram cell dendrites (Figure S4). In contrast to CS+, there was no significant change in the evoked dendritic activity in response to CS- in either engram cells (baseline, 0.04 ± 0.01 Hz vs CS-, 0.03 ± 0.01; p = 0.54; Figure 2c, e) or non-engram cells (baseline, 0.04 ± 0.01 Hz vs CS-, 0.04 ± 0.01; p = 0.48; Figure 2d, f).

**Figure 2.**
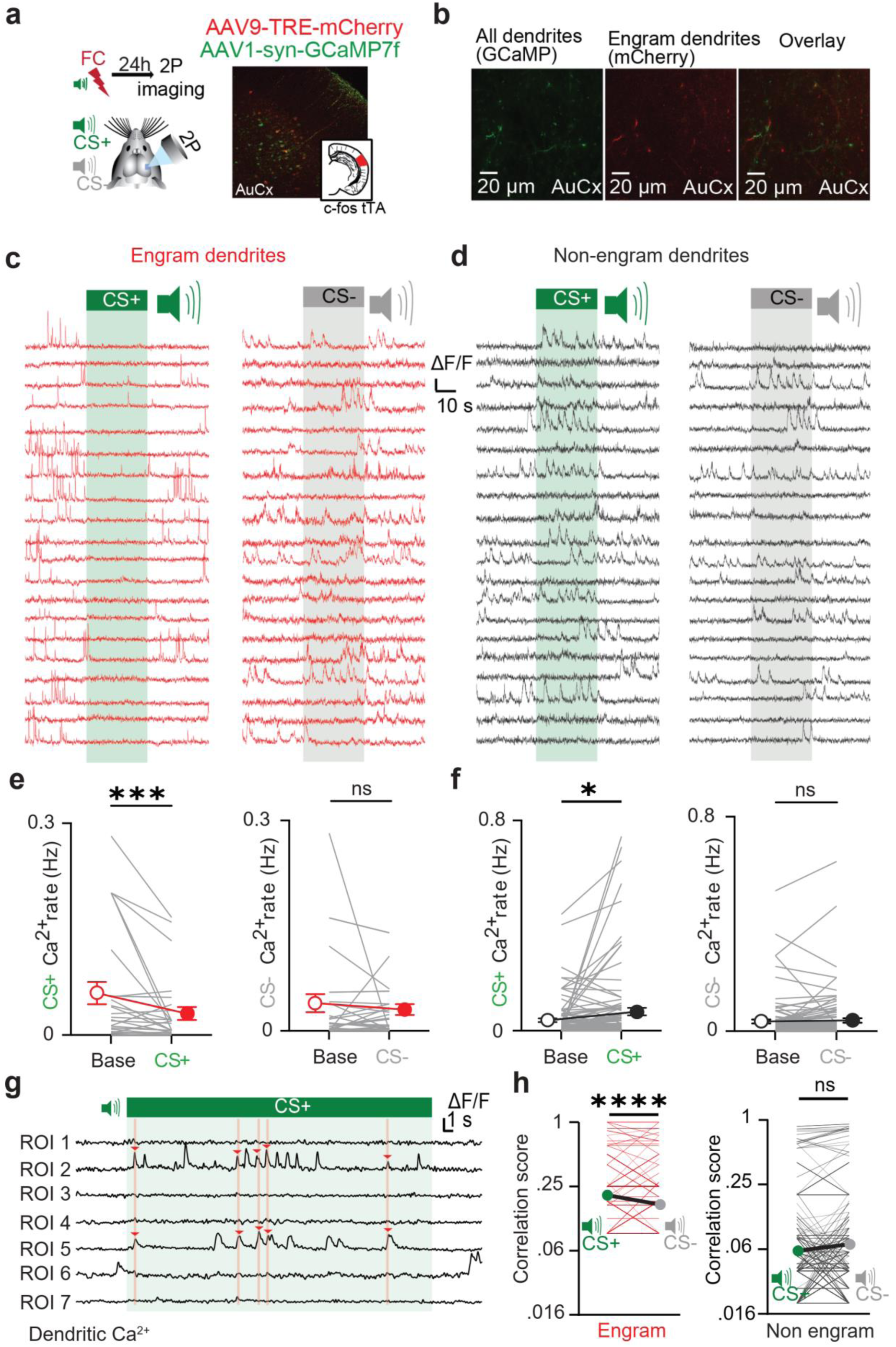
Low and synchronized activity of engram cells tuft dendrites during memory recall. **a,** Experimental design and representative pictures. Engram cells were labelled with mCherry (AAV_9_-TRE-mCherry) and a calcium indicator was expressed in all neurons (AAV_1_-syn-GCaMP7f). 24h after conditioning, awake mice were head fixed under a 2 photons microscope to image the activity of their tuft dendrites during exposure to the CS+ and CS-. Picture of a fixed brain slice showing the expression of engram cells (red) and GCaMP7f (green) in the auditory cortex. **b,** 2 photons microscope pictures of GCaMP7f (left) and mCherry (middle) signals and their overlay (right) at the tuft dendrites. **c-d,** Example traces one EC (c) and one NEC (d) dendrite during CS+ and CS- exposure. Each trace represents a different occurrence of the tone. **e-f,** Quantification of the rate of calcium events of EC (E; n = 27 from 7 mice) and NEC (F; n = 106 from 7 mice) tuft dendrites during CS+ and CS- exposures. A significant decrease of EC tuft dendrites activity, and significant increase of activity of NEC tuft dendrites during the CS+ specifically. **g,** Methods for the detection of synchronous events. Each trace represents a different NEC dendrite from the same mouse during one CS+. If events peaked during the same time window (pink bars, 133 ms duration) they were considered coincidental (red arrows). **h,** Average correlation score for EC (left, red) and NEC (right, grey) dendrites during CS+ and CS- exposures. EC dendrites show an increased synchronous activity specifically during CS+ exposure. Data are represented on a logarithmic scale. Data are represented as mean ± SEM and significance was assessed with a two-tailed paired Wilcoxon for e and f (*: p < 0.05, **: p < 0.001), and a two-tailed unpaired Mann-Whitney test for h (****: p < 0.0001).

Given the importance of synchronized activity in sensory perception^22^, we next evaluated the synchronization of dendritic activity during tone exposure. Here, calcium transients in engram and non-engram dendrites within the same trial were assigned a correlation score according to how many other dendrites had evoked activity at the same time (Figure 2g; see Methods). Engram cell dendrites had significantly greater correlated activity during CS+ (0.20 ± 0.01) compared to CS- (0.17 ± 0.004; p < 0.0001; Figure 2h). This correlation score was also significantly higher compared to a randomly shuffled dataset, confirming the result (Figure S4; p = 0.026). In contrast, the dendrites of non-engram cells had no significant difference in the correlation score in response to CS+ compared to CS- (0.06 ± 0.002 vs 0.07 ± 0.002; p = 0.3; Figure 2g) nor with a randomly shuffled dataset (p = 0.16; Figure S4). Taken together, these results show that engram cells have decreased, but synchronized, dendritic activity during fear memory recall.

### Local engram network in the auditory cortex

A possible explanation for the increased synchronization of engram dendrites is the presence of an engram network in the auditory cortex. To test this hypothesis, we labelled engram cells with AAV_9_-TRE-EYFP-ChR2 and performed *ex vivo* patch-clamp voltage recordings 24 h after fear conditioning. Engram cells were identified both visually (YFP) and functionally (photo-response to a 500 ms light pulse; Figure 3a-b). To assess connectivity, engram cells were photoactivated with a single brief 15 ms pulse of blue LED over the slice surface. To ensure all recorded responses were mono-synaptic, the sodium and potassium channels antagonists (TTX and 4AP respectively) were included in the bath throughout experiments^44^ (Figure 3c). LED activation not only directly photo-activates engram cells, it also activates the pre-synaptic axon terminals. Blocking synaptic transmission with bath application of CNQX (AMPA antagonist, 10 µM) and APV (NMDA antagonist, 100 µM) decreased the amplitude of the ChR2-evoked response (by 4.85 ± 1.02 %; p < 0.001; n = 13 from 11 mice; Figure 3d), suggesting the recorded evoked response included synaptic inputs from other engram cells. These results suggest that, similar to other brain regions^8,11,45,46^, engram cells are synaptically connected with surrounding engram cells, forming a local network in the auditory cortex. This decrease in the photoactivated response was not due to the waiting time during drug application, as the light response measured again 10 min later in a subset of cells showed no significant decrease due to time alone (data not shown). Subtracting the LED-evoked potential following the synaptic block enabled the isolation of the ChR2-evoked EPSP in engram cells (Figure 3e). Following subtraction, synaptic inputs were observed in 84.6 % of the recorded EC (11/13 cells), whereas only 14.7% of recorded NEC (5/32 cells) presented a light response (Figure 3f). Compared to non-engram cells, the LED-evoked potential in engram cells was significantly greater in amplitude (0.36 ± 0.08 (engram cells, n = 13 from 11 mice) vs 0.02 ± 0.02 mV (non-engram cells, n = 32 from 16 mice); p < 0.0001; Figure 3g, h). Combined, the increased number of responsive engram cells and the larger synaptic response during engram cell photoactivation suggests that engram cells preferentially target other engram cells compared to non-engram cells. Taken together, these results suggest the existence of an intra-connected engram network in the auditory cortex.

**Figure 3.**
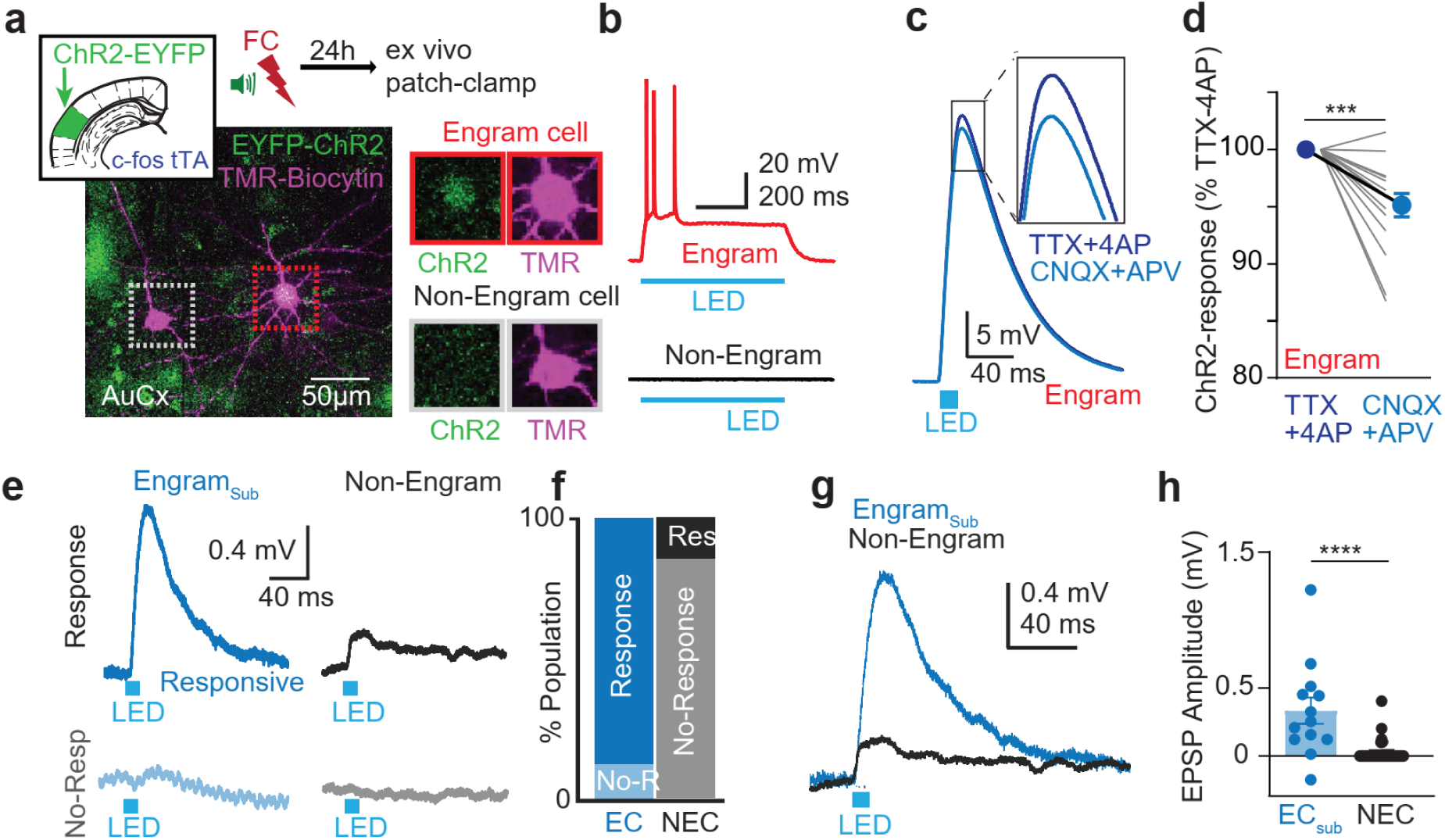
Local engram network in the auditory cortex. **a,** An AAV_9_-TRE-EYFP-ChR2 was injected in the auditory cortex to label engram cells as described previously. Confocal picture showing a recorded EC (red square) and a neighbouring NEC (grey square) filled with biocytin during recording. **b,** Example traces showing the voltage response of an EC (top, red) and a NEC (bottom, grey) to a 500 ms pulse of blue light. A depolarisation lasting for the entire pulse confirmed intrinsic ChR2 expression in EC. **c,** Example trace showing the voltage response of an EC to a 15 ms pulse of blue light. EC to EC connectivity was evaluated by measuring the decrease of the light response after glutamatergic blockade with CNQX (10 μM) and APV (100 μM). Monosynaptic inputs were insured by adding TTX (1μM) and 4AP (5oo μM) to the bath. **d,** Amplitude of the voltage response at baseline (TTX+4AP) and under blockade of glutamatergic transmission (CNQX+APV). Values were normalized on the baseline response (TTX+4AP). A significant decrease of the response was observed in EC when glutamatergic transmission was blocked (n = 13 from 11 mice). **e,** Example of a subtraction trace from an EC showing the voltage response that was blocked under CNQX+APV (dark blue, top left), and a raw voltage response to light stimulation in a NEC (dark grey, top right). Bottom traces show example traces when no synaptic input in response of blue light was detected in an EC (left, light blue) and an NEC (right, light grey). **f,** Proportions of responsive (dark colours) and non-responsive (light colours) neurons in the engram population (blue) and the non-engram population (grey). A higher proportion of EC received synaptic inputs from other EC compared to NEC. **g,** Overlay of example traces of responsive neurons shown in (f). **h,** Quantification of the amplitude of the synaptic responses for EC (n = 13 from 11 mice) and NEC (n = 32 from 16 mice). EC have a significantly greater amplitude of synaptic inputs compared to NEC. Data are represented as mean ± SEM and significance was assessed with a two-tailed paired Wilcoxon for (D) (***: p < 0.001), and a two-tailed unpaired Mann-Whitney test for (H) (****: p < 0.0001).

### Decreased dendritic excitability in engram cells is due to increased Ih

To understand the dendritic mechanisms underlying the pattern of activity observed in engram cells, we used *ex vivo* patch-clamp recordings combined with extracellular stimulation of tuft dendrites. Engram cells were labelled with an AAV_9_-TRE-mCherry and 24 h later the brain was sliced for patch-clamp recordings. Neurons were filled with a fluorophore (TMR-biocytin) via the patch pipette to visualize the dendritic arbour, and an extracellular stimulation theta electrode was placed in proximity (within 5 µm) to a tuft dendrite under visual guidance (Figure 4a). Supralinear voltage responses were evoked in both engram (Figure 4b) and non-engram (Figure 4c) cells in response to double-pulse extracellular stimulation (0.1ms, 100 Hz) of increasing intensity. The resulting supralinear voltage response evoked in engram cells was similar in amplitude to non-engram cells (Figure 4e), but had a significantly reduced half-width (38.77 ± 1.95 ms (engram cells, n = 4 from 4 mice) vs 93.85 ± 14.29 (non-engram cells, n = 4 from 4 mice); p = 0.029; Figure 4e) and decay time (55.08 ± 3.04 ms vs 97.02 ± 12.15; p = 0.029; Figure 4G). This characteristic makes engram tuft dendrites ideally suited for coincidence detection^38^, in line with their synchronous activity observed *in vivo*. In comparison with non-engram cells, engram cells had increased voltage sag in response to a hyperpolarizing current injection (13.53 ± 0.97 % and 9.32 ± 1.16 of steady state; p = 0.009; n = 7 from 7 mice (engram cells) and n = 9 from 9 mice (non-engram cells); Figure 4h), illustrating engram cells had larger Ih currents. Since Ih is known to reduce input summation^47^ and shorten EPSPs by speeding the decay of the response^48^, we further investigated the influence of greater Ih in engram dendrites on the integration of synaptic input. In response to a train (5 pulses, 50 Hz) of extracellular stimulation inputs targeted to tuft dendrites, both engram and non-engram cells evoked a voltage response which was increased by blocking Ih with bath application of ZD7288 (20 µM; Figure 4i, j). Overall, Ih block resulted in a greater increase in the evoked voltage response in engram cells (244.40 ± 58.71 %, n = 9 from 9 mice) compared to non-engram cells (183.10 ± 23.40 %; n = 7 from 7 mice; p = 0.011; Figure 4k). Furthermore, the half-width of the evoked voltage response was significantly increased during Ih block in engram cells (aCSF, 36.91 ± 6.06 ms vs ZD7288, 82.08 ± 6.07 ms; p = 0.0001) but there was no effect on non-engram cells (aCSF, 48.33 ± 9.19 ms vs ZD7288 61.23 ± 11.05 ms; p = 0.12; Figure 4i). Taken together, these results illustrate that dendritic Ih specifically dampens the voltage response in engram cells, suggesting a crucial role in the low and synchronized pattern of activity observed *in vivo*.

**Figure 4.**
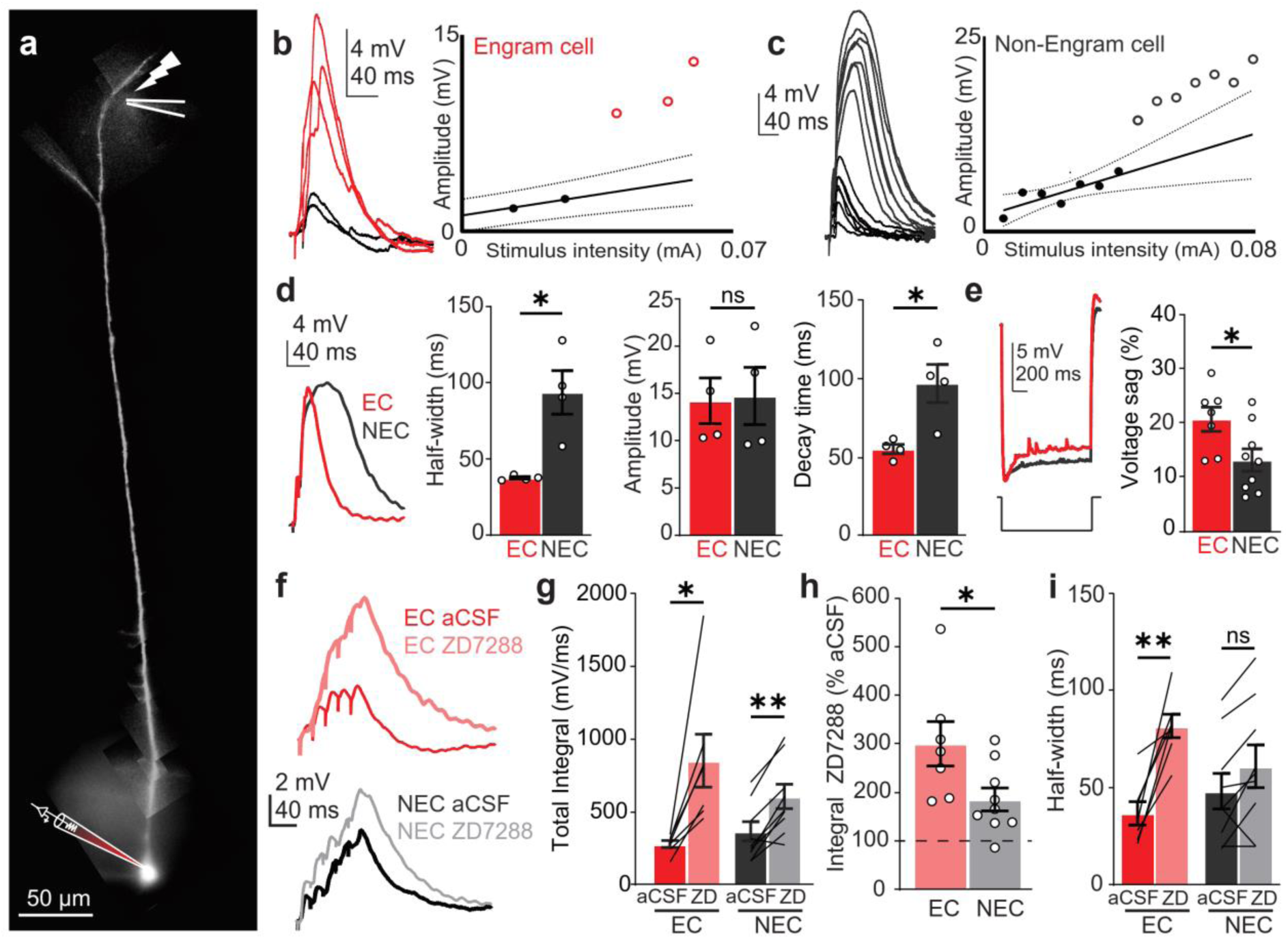
Increased Ih in EC decreases excitability of their tuft dendrites and sharpens the voltage response to synaptic inputs. **a,** Example picture of a recorded neuron filled with biocytin-TMR allowing the visualisation of the dendritic arbour, allowing to target tuft dendrites with the extracellular stimulation. The stimulation electrode is highlighted at the top. **b-c,** Example traces and corresponding input/output (I/O) scatter plots of supralinear events evoked at the tuft dendrites in an EC (left) and a NEC (right). Plain lines show the linear fit of the early part of the I/O curve, and dashed lines the 95% confidence interval of this fit. Response outside of the confidence interval are considered supralinear events. Supralinear events are represented with empty coloured circles, and their corresponding traces are coloured (EC: red, b; NEC: grey, c). **d,** Left: overlay of example traces of a supralinear events evoked in EC (red) and NEC (dark grey). Right: bar graphs showing the quantification of the half-width, amplitude and decay of supralinear events, revealing a significantly shorter half-width and decay time in EC for events of the same amplitude (n = 4 from 4 mice (EC and NEC)). **e,** Example traces (left) and quantification (right) of the voltage sag in response to a hyperpolarizing step of current injection, indicative of Ih currents. Results reveal significantly increased Ih in EC (EC: n = 7 from 7 mice; NEC: n = 9 from 9 mice). **f,** Example traces of the response to a high frequency train of stimulation (50Hz, 5 pulses) in EC (top) and NEC (bottom) without (dark traces) and with (light traces) the Ih antagonist ZD7288. **g,** Bar graph showing the integral of the voltage response to the 5 pulses for EC and NEC under baseline (aCSF) and Ih block with ZD7288 (ZD). A significant increase is observed in both populations. **h,** Bar graph showing the integral of the response under ZD7288 as a percentage of the baseline response (aCSF). The dashed line represents 100% (response under aCSF). The increase of the response was significantly larger in EC (n = 7 from 7 mice) compared to NEC (n = 9 from 9 mice). **i,** Quantification of the half-width of the response to the 5 pulses stimulation at baseline and under ZD7288. A significant increase of the half-width by ZD7288 was observed only in EC. Data are represented as mean ± SEM and significance was assessed with a two-tailed unpaired Mann-Whitney test for d, e and h (*: p < 0.05), and a two-tailed paired Wilcoxon test for g and h (*: p < 0.05, **: p < 0.01).

### Learning-induced Ih transient upregulation decreases engram cells excitability

To test whether increased Ih in engram cells was learning-induced plasticity, we performed *ex vivo* patch-clamp recordings either 24 h or 1 week after conditioning while Ih currents were blocked by bath application of the antagonist ZD7288 (Figure 5a-c). 24 h following learning, blocking Ih increased the excitability of engram cells, significantly increasing the evoked firing rate during current step injection (from 30 to 230 pA; Figure 5D and E), leading to a significantly increased average firing rate (9.20 ± 0.70 Hz (aCSF, n = 46 from 30 mice) vs 14.19 ± 1.50 (ZD, n = 20 from 13 mice); p < 0.001; Figure 5f). Ih block also increased the membrane resistance (68.71 ± 2.77 MΩ (aCSF) vs 106.10 ± 5.44 (ZD); p < 0.0001) and decreased the rheobase (114.90 ± 5.90 pA (aCSF) vs 81.58 ± 13.10 (ZD); p < 0.05; Figure S5). In contrast, block of Ih current had a reduced impact on non-engram cells, only increasing firing rate in response to a smaller range of current injections (Figure 5g, h). In addition, Ih blockage had not effect on the overall average firing rate (10.3 ± 0.7 Hz (aCSF, n = 25 from 22 mice) vs 11.8 ± 1.5 (ZD, n = 12 from 10 mice); p = 0.61; Figure 5i) and rheobase (94.80 ± 7.77 pA (aCSF) vs 71.67 ± 14.24 (ZD); p = 0.29; Figure S5). Since blocking Ih resulted in similar intrinsic properties in engram and non-engram cells, our data suggest that Ih is sufficient to explain the decreased excitability in engram cells recorded 24 h following fear learning. One week following fear learning, there was no significant difference in the voltage sag (11.9 ± 1.9 % (engram cells, n = 14 cells from 6 mice) vs 11.2 ± 2.2 (non-engram cells, n = 13 cells from 6 mice; p = 0.7; Figure 5k) or overall cellular properties between engram and non-engram cells (Figure S6). At this time points, Ih blockage had no effect on the evoked firing rate in engram cells (10.87 ± 2.04 (aCSF, n = 14 from 6 mice) vs 11.13 ± 2.26 Hz (ZD, n = 9 from 4 mice); p = 0.91; Figure 5m-o) or non-engram cells (9.84 ± 1.63 (aCSF, n = 13 cells from 6 mice) vs 11.68 ± 2.08 Hz (ZD, n = 8 from 4 mice); p = 0.80; Figure 5p-r), nor on the rheobase and membrane resistance (Figure S6). Taken together, these results suggest a transient learning-induced up-regulation of Ih in engram cells that temporarily decrease their somatic and dendritic excitability following fear learning.

**Figure 5.**
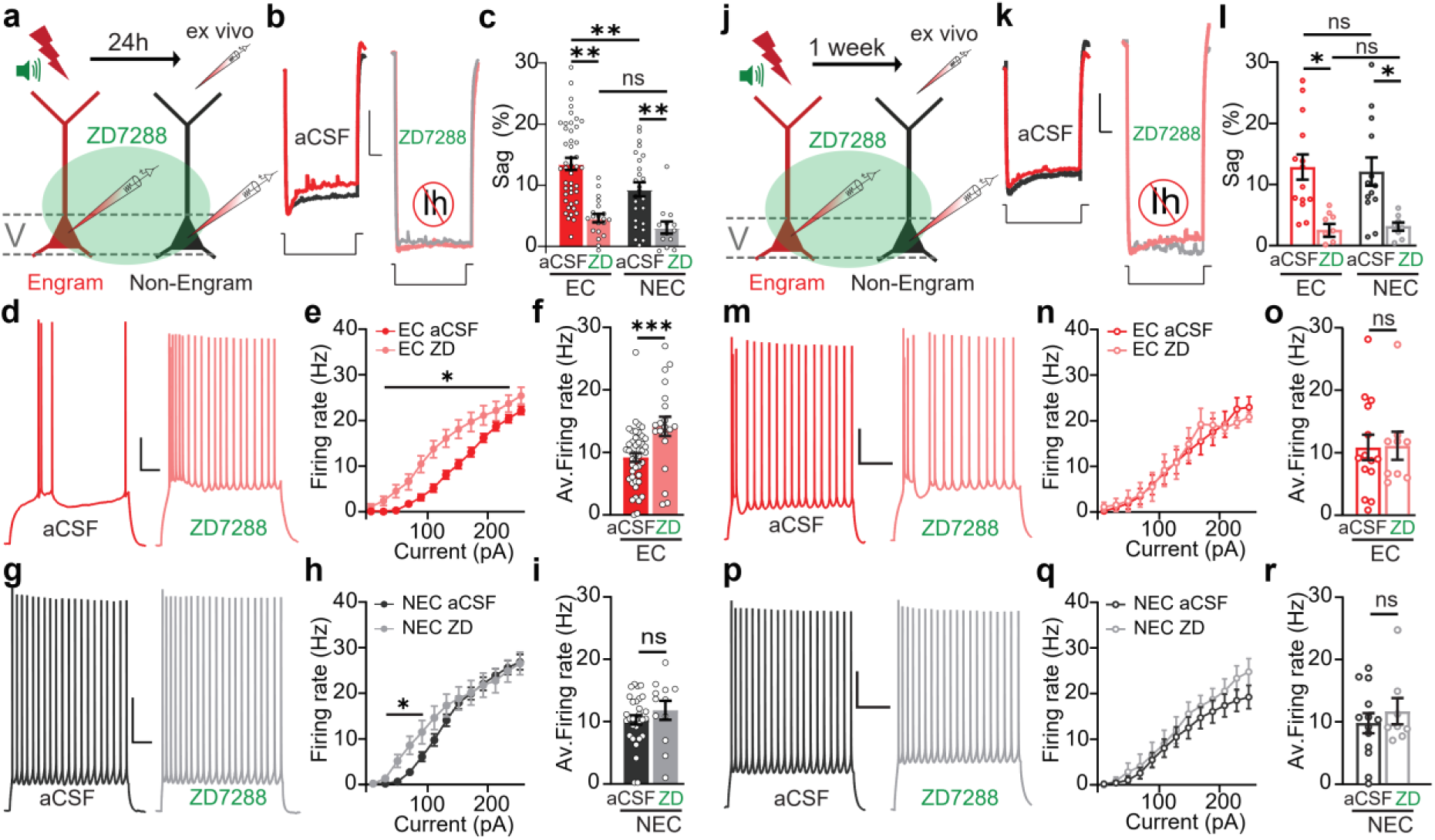
No difference in Ih nor excitability at later time points reveal transient learning-induced intrinsic plasticity in EC. **a,** *Ex vivo* patch-clamp recordings of layer 5 EC and NEC were performed 24h after conditioning. Intrinsic properties were measured by injecting current steps of increasing intensities under control condition (aCSF) and under Ih blockade by bath application of ZD7288. **b-c,** Example traces (b) and quantification (c) of the voltage sag in response to a hyperpolarizing step of current injection, indicative of Ih currents under aCSF and ZD7288. An increased voltage sag is observed in EC, and its efficient blockade by ZD7288. Scale bars represent 4 mV/200 ms. **d,** Example traces of action potentials firing in EC under aCSF (left, red) and ZD7288 (right, pink) in response to a 130 pA current injection. Scale bars represent 20 mV/200 ms. **e,** Action potential firing as a function of injected current in EC under aCSF (red, n = 46 from 30 mice) and ZD7288 (pink, n = 20 from 13 mice). **f,** Average firing rate of EC over the 10-250 pA range of current injections, under aCSF and ZD7288. A significant increase under ZD7288 is observed. **g,** Example traces of action potentials firing in NEC under aCSF (left, dark grey) and ZD7288 (right, light grey) in response to a 130 pA current injection. Scale bars represent 20 mV/200 ms. **h,** Action potential firing as a function of injected current in NEC under aCSF (dark grey, n = 25 from 22 mice) and ZD7288 (light grey, n = 12 from 10 mice). **i,** Average firing rate of NEC over the 10-250 pA range of current injections, under aCSF and ZD7288. No significant increase under ZD7288 is observed. **j,** *Ex vivo* patch-clamp recordings of layer 5 EC and NEC were performed 1 week after conditioning. Intrinsic properties were measured by injecting current steps of increasing intensities under control condition (aCSF) and under Ih blockade by bath application of ZD7288. **k-l,** Example traces (k) and quantification (l) of the voltage sag in response to a hyperpolarizing step of current injection, indicative of Ih currents under aCSF and ZD7288. No difference between EC and NEC was observed at that time point. Scale bars represent 4 mV/200 ms. **m,** Example traces of action potentials firing from 1 EC 1 week following conditioning under aCSF (left, red) and ZD7288 (right, pink) in response to a 170 pA current injection. Scale bars represent 20 mV/200 ms. **n,** Action potential firing as a function of injected current in EC 1 week following conditioning under aCSF (red, n = 14 from 6 mice) and ZD7288 (pink, n = 9 from 4 mice). **o,** Average firing rate of EC 1 week following conditioning over the 10-250 pA range of current injections, under aCSF and ZD7288. No significant effect of ZD7288 is observed. **p,** Example traces of action potentials firing from 1 NEC 1 week following conditioning under aCSF (left, dark grey) and ZD7288 (right, light grey) in response to a 170 pA current injection. Scale bars represent 20 mV/200 ms. **q,** Action potential firing as a function of injected current in NEC 1 week following conditioning under aCSF (dark grey, n = 13 from 6 mice) and ZD7288 (light grey, n = 8 from 4 mice). **r,** Average firing rate of NEC 1 week following conditioning over the 10-250 pA range of current injections, under aCSF and ZD7288. No significant effect of ZD7288 is observed. Data are represented as mean ± SEM and significance was assessed with two-tailed unpaired t-tests (*: p < 0.05, **: p < 0.01, ***: p<0.001)

## Discussion

We identified a population of engram cells within layer 5 (L5) of the auditory cortex with decreased excitability and increased connectivity. These L5 pyramidal neurons were classified as engram cells as 1) they were activated by learning and recall, 2) their optogenetic reactivation can induced specific learned behaviour and 3) they showed specific *in vivo* activity and plasticity as a result of learning. As the site of feedback information crucial for learning and recall^23,25,27^, the apical tuft dendrites of engram cells had low, but synchronized activity during memory recall following fear conditioning. Our results show that plasticity of Ih, a crucial current in dendritic integration, primarily drove the differences in excitability between engram and non-engram cells, which reduced the excitability of tuft dendrites and decreased the somatic output. This engram-specific plasticity of dendritic integration declined a week after learning, suggesting learning-induced transient Ih plasticity in cortical engram cells. Together, these findings suggest learning can result in the formation of a cortical engram with distinct circuit properties and somatic and dendritic plasticity.

### Decreased and synchronized activity of engram cells tuft dendrites

As a target of feedback input, cortical layer 1 has been identified as critical for memory processes, especially learning^24,25^. In the current study, we found that the tuft dendrites of engram cells are active in a low but synchronous manner during memory recall. The decreased activity challenges the hypothesis that engram cells necessarily are more active than non-engram cells during memory recall. As L5 pyramidal cells are the main output of the cortex, these engram cells could be central to connect and synchronize the auditory cortex with the brain wide engram. This is supported by the biophysical and synaptic properties of engram cells revealed by our *ex vivo* recordings, which illustrated that engram cells have a briefer suprathreshold dendritic voltage response and greater burst firing, which are important in the synchronization of neuronal activity^38,49^. Moreover, our results show that these parameters are impacted by Ih conductance, which is increased in engram cells. Since Ih is a critical current in dendritic integration which decreases dendritic excitability and shortens synaptic response^38,47^, it is likely to strongly influence the low and synchronized activity of engram cells during memory recall. This decreased excitability due to increased Ih in engram cells may lead to sparse coding within the engram cell population, a type of information coding previously observed in sensory cortices^50,51^ that is increased after learning^52^.

### Ih-dependent intrinsic plasticity

It has been proposed that the excitability of neurons at the time of learning determines their allocation to the engram, with the most excitable cells becoming engram cells^7,8,53^. In the current study, engram cells had decreased firing 24 h following conditioning, suggesting that this decreased excitability would be due to learning-induced plasticity in engram cells. This difference in excitability in engram cells was abolished one week after fear conditioning. This suggests a transient learning-induced upregulation of Ih in engram cells, temporarily reducing their global excitability, both at the dendrites and soma. Learning- and burst firing-induced plasticity of Ih have been shown in previous studies^39,40,48,54^, and may be at the heart of the changes in neural encoding following other learnings, in addition to fear learning.

### General conclusions

The specific role of dendrites in memory formation is seldom investigated, and this study illustrates changes in dendritic properties may underlie the changes in cellular properties required for memory formation. Overall, this study highlights Ih plasticity in engram cells as a potential mechanism which drives the transient decrease in both the somatic and dendritic excitability. By influencing input integration, and most likely the activity of the downstream engram network, dendritic Ih plasticity plays a central role in memory processes within single cortical neurons which may be central to connect and synchronize the auditory cortex with the brain wide engram.

## Methods

All experiments were conducted in accordance with the Code of Practice for the Care and Use of Animals for Scientific Purposes (National Health and Medical Research Council, Australia). The guidelines were given by the veterinary office and approved by the Animal Ethics Committees of The Florey Institute of Neuroscience and Mental Health, University of Melbourne, and of the Australian National University.

### Subjects

All animals were 6-14 weeks old male and female cfos-tTA/cfos-EGFP mice (Jax reference 008344). They were housed with constant access to fresh water and doxycycline-containing diet (40 mg/kg, SpecialtyFeeds; SF16-087) on a 12/12 light/dark cycle.

### Engram cell labelling strategy

Engram cells were labelled using an activity-based strategy^3,43^. A virus expressing a transgene under the control of a tetracycline-responsive element (TRE) promoter was injected in transgenic mice expressing the tetracycline transactivator (tTA) under the c-fos promoter. To restrict the expression to the learning period, animals were constantly fed with a doxycycline-containing diet (40 mg/kg) from at least 10 days before virus injection. This diet was replaced by a doxycycline-free diet only during the 2 days preceding learning to allow for the labelling of engram cells.

### Adeno-associated viruses

The AAV vectors used for viral production for engram labelling were pAAV-TRE-ChR2-EYFP, pAAV-TRE-EYFP and pAAV-TRE-mCherry. Viruses were made by Vigene Bioscience with DNA minipreps custom made. Viruses were injected with the following titers (in genome copy per mL): 9.98×10^13^ (AAV_9_-TRE-EYFP), 9.03×10^13^ (AAV_9_-TRE-EYFP-ChR2) and 2.54×10^13^ (AAV_9_-TRE-mCherry). During in vivo calcium imaging, AAV_1_-hsyn-GcaMP7f was used (Addgene reference #104488)

### Stereotactic virus injection

Surgeries were performed using a stereotaxic apparatus (Narishige) under isoflurane anaesthesia (1-2.5% in 0.75-1 L/mn O2), and body temperature was maintained at 36-37 °C using a heat pad. Eye ointment was applied to prevent dehydration and meloxicam (Ilium) was intraperitoneally injected at the end of surgery. The head was shaved and the skin disinfected with betadine, before cutting the skin to expose the skull. Craniotomies were performed using a 0.4 mm diameter drill (Komet dental) above the auditory cortex (from bregma: + 2.4 mm AP, ± 4.2 to 4.5 mm ML; - 0.9 mm DV from brain surface). A pulled glass pipette (Drummond) was inserted in the brain. Viruses were injected (60-100 nL/mn) using a manual injector (Narishige) 1 minute after reaching the injection coordinates. Pipettes were withdrawn 5 minutes after the injection was done, and the skin was closed with stitches. Behavioral procedures started 1-2 weeks after injection, or at least 3 weeks after injection for *in vivo* calcium imaging.

### Tone fear conditioning and recall

#### Apparatus

All experiments were conducted in Med Associates fear conditioning chambers (22cm x 33cm x 25cm). Electric shocks were delivered through a grid floor. Different contexts were used for conditioning (context A: infrared light, grid floor, black triangular walls, benzaldehyde smell (Sigma-Aldricht, 0.25%)), natural recall (context B: white light, white circular walls, white smooth floor, acetic acid (Sigma-Aldricht, 1%)), and optogenetic sessions (context C: no ceiling, black and white striped circular walls and floor, ethanol 70%).

#### Habituation, conditioning and recall

Mice were habituated to handling and transport 10 minutes per day for 3 days before experiments started. After habituation, animals were placed on a doxycycline-free diet in a quiet room for 48h before being conditioned, then put back in a quiet room for 24h on a doxycycline diet (40 mg/kg). Conditioning consisted in a 10.5 minute session with 5 CS+/US pairings starting after 2 minutes and separated by 1 minute each. The CS+ was a 30 sec succession of 500 ms sweeps (70 dB) interleaved with 500 ms of silence. Each sweep was composed by a train of 5 pure tones of increasing frequencies (90 ms each, 5 KHz to 15 KHz with 2.5 KHz steps) separated by 10 ms. The US was an electric footshock (2 sec, 0.6 mA) that co-terminated with the CS+ (see Figure S1 for a visual description of the CSs). Natural recall was conducted 24h after conditioning in context B, and 5 CS+ were played without the US being delivered. For all behavioral experiments, animals were brought to the context in a transfer cage, and the chambers were cleaned with ethanol (80%) between each animal. Memory was assessed by manually quantifying freezing behavior for each epoch of 3 seconds during the recall session.

### Optogenetic activation of engram cells

#### Experimental groups

One group was injected with the AAV_9_-TRE-ChR2-EYFP and conditioned as described above (Shock group). To control for the effect of the reactivation of the auditory cortex when no fear was involved, one group was injected with the same virus but only exposed to the CS+ without footshock (No Shock group). To control for any fear generalization, another group was injected with the AAV_9_-TRE-EYFP and conditioned following the same procedure (No ChR2 group).

#### Surgical procedure

Virus injection was performed with an angle at adjusted coordinates (from bregma: + 2.4 mm AP, ± 4.5 to 4.8 mm ML; - 0.45 mm DV from brain surface). Once the skull drilled, the head of the animal was tilted with a 45° angle and the pipette was inserted in the brain for virus injection. Once the injection done, an optic fiber cannula (400 µm diameter, Thorlabs) was placed in contact with the brain surface and fixed with metabond (Sun Medical, Japan). The injection and cannula implantation were done for one hemisphere before switching to the other one. After both cannulas were implanted, small parts of pipette cone tips were placed around as protection, and everything was cemented together in dental cement. Dental cement was mixed with ink to reduce light scattering during optogenetic sessions.

#### Handling and habituation

Mice were habituated to handling and transport for 5-15 minutes per day for 3 days before habituation to the patch cord and optogenetic context started. During habituation, animals were plugged to the patch cord (400 µm diameter, thorlabs) connected to a rotary joint (Doric) and an LED (Doric). Animals were left to explore the context for 15 minutes each day for 4 consecutive days. Blue light (470 nm, 5-15 mW/mm) was delivered for 6 epochs lasting 1 minute and separated by 1 minute, starting after 2 minutes. On days 1 and 3, light was delivered at 20Hz (15 ms pulses), and on days 2 and 4 at 4 Hz (15 ms pulses). On days 3 and 4, animals were exposed to 3 x 2 mn of CS-, co-starting with the 1st, 3rd and 5th light epochs. The CS- was a 30 sec succession of 500 ms sweeps (70 dB) interleaved with 500 ms of silence. Each sweep was a broadband noise of the same frequency range than the CS+ (5 to 15 KHz). After habituation, engram cells were labelled as previously described.

#### Natural Recalls

24h after conditioning (natural recall 1) and 24h after the last optogenetic session (natural recall 2), animals were placed in context B and exposed to either the CS+ or CS-. Memory to the CS+ and CS- was tested during 2 different sessions separated by at least 2 hours on the same day. One recall session consisted in 3 exposures to the CS interleaved by 1 minute and starting 2 minutes after the animal entered the context.

#### Optogenetic reactivation

One optogenetic session was conducted each day for 8 days and from day 2 post-conditioning. Light intensity was measured before each session started. 4 different conditions were tested twice each over the experiment (between day 1 and 4, and a second time between days 5 and 8). Each condition consisted in the combination of a frequency of reactivation (20 Hz or 4 Hz) with or without the CS- being played. Results shown in figure 1H are averaged over 2 days of the same condition (days 4 and 8, 4Hz light + CS-).

### Histology

#### Tissue fixation and conservation

Within 2 weeks after engram cells labelling, mice were deeply anaesthetized with a lethal dose of urethane (20%) and transcardially perfused with phosphate buffer (PB 0.1 M) followed by 4% paraformaldehyde (PFA) diluted in PB. Brains were placed in PFA 4% overnight then in PB 0.1 M and kept at 4°C until slicing. For c-Fos quantification, perfusions were performed 1h to 1h15 after the recall session.

#### Immunostainings

Coronal slices (50 µm thick) were performed in PB 0.1 M using a vibratome (Leica), collected in PB 0.1 M and kept at 4°C. The immunostaining procedure started with a saturation step (1h at room temperature in phosphate buffer saline with 0.3 % triton X-100 (PBST)), with 10 % bovine serum albumin (BSA, sigma-aldricht) followed by incubation in the primary antibodies solution constantly shaking overnight at 4°C. After 3 x 10 min washes in PB 0.1 M shaking at room temperature, slices were incubated in the secondary antibodies solution for 3h, constantly shaking at room temperature. All antibodies were diluted in PBST with 3 % BSA: 1:1000 for primary antibodies (chicken anti-GFP (Abcam, ab13970), rabbit anti-c-Fos (synaptic system, 226 003), rat anti-mCherry (Invitrogen, M11217)), and 1:500 for secondary antibodies (goat anti-chicken Alexa 488 (invitrogen, a-11039), donkey anti-rabbit Alexa 488 (Invitrogen, a-21206), goat anti-rabbit Alexa 568 (Invitrogen, a-11011), goat anti-rat Alexa 555 (Invitrogen, a-21434)). Slices were washed 3 x 10 mn in PB 0.1M shaking at room temperature before being mounted in mounting medium (Fluorophore with DAPI, Sigma-Aldricht) for further imaging.

#### Confocal imaging

Fluorescent images of the auditory cortex were obtained with a confocal microscope (Leica SP8). For further quantification, Z-stacks (3 µm steps) of the entire slice were taken with an 40x oil objective. For injection site and virus expression verification, a single image was taken with a 20x air objective.

#### Cell counting

All quantifications were performed on at least 4 slices per animal, and all counting were performed manually using the CellCounter plugin (ImageJ). Engram cells labelling was evaluated by counting the number of mCherry or EYFP expressing cells. The number of DAPI cells per surface unit in the auditory cortex was evaluated in 3 animals by counting DAPI nuclei from one focal plane. This estimated value was used to express the number of engram cells per DAPI cells in all animals. Recall-induced engram reactivation was evaluated by quantifying the co-expression of c-Fos with mCherry or EYFP. Superficial and deep layers were counted separately for all analysis.

### Calcium imaging

#### Surgical procedure

Virus injection was performed in the left hemisphere as described in the stereotactic injection paragraph. A cocktail of AAV_1_-hsyn-GcaMP7f (3:4) and AAV_9_-TRE-mCherry (1:4) was injected in the auditory cortex (2.4mm posterior to bregma, 4.2 to 4.5mm lateral of midline, 0.9mm deep from brain surface). A 3mm craniotomy was performed above the injection site using a biopsy punch (Kai Medical), and a circular coverslip was glued on top. A headbar was placed on the skull on the right hemisphere and everything was fixed in dental cement.

#### Habituation

Mice were habituated to head-fixation over at least three days, with increasing durations over the course of habituation (a few seconds to a few minutes).

#### Imaging

Calcium imaging was performed using a two-photon microscope (Sutter Instruments, Moveable Objective Microscope) with a femtosecond tuneable laser (SpectraPhysics Maitai DeepSee) controlled with the ScanImage software^63^. Photons emitted from jGCamP7f or mCherry were captured following separation into green and red light with a dichroic filter (565dcxr, Chroma Technology) before being detected with GaAsp photomultiplier tubes (Hamamatsu).

Mice were head-fixed and a speaker taken from one of the fear conditioning boxes (Med Associate) was placed close (∼5 cm) to the animal’s head. An imaging session consisted of interleaved CS+ and CS- with an inter-trial interval of randomized duration (30 to 60 seconds). Each CS was played 20 times at an intensity of 70 dB.

On the day of imaging, engram cells were identified at 1040 nm to identify mCherry labelling. Each engram cell’s dendritic tree was surveyed at 1040nm to build a picture of its branch morphology. The green channel with excitation at 840nm and 940nm were used to build a picture of all GCaMP labelled neurons. This process required many changes in location, zoom, and wavelength, all observed by at least 2 experimenters to confirm dual labelling. The Layer 5 origin of the engrams were confirmed by tracking the apical trunk down past maximum visible distance (∼400-500 µm). The field of view (FOV) selected focused on tuft dendrites of as many engram neurons as possible (two or three), positioned to also include as many non-engram dendrites as possible. Full confirmation of dual labelled cells after the session was performed through manual alignment of a red channel 1040nm excitation z-stack and a green channel 840nm z-stack, both locally to the FOV using high spatial resolution and across the entire dendritic tree at lower spatial resolution.

GCaMP fluorescence activity was acquired on the green channel at 940nm excitation. Motion correction was performed using a cross-correlation algorithm. Regions of interest (ROIs) were drawn around individual dendritic branches of L5 neurons (identified with the z-stacks), using ImageJ’s freehand tool over a mean fluorescence image. Fluorescence extracted from the ROIs had *ΔF/F* calculated for both the ROI and a background mask (defined by dilating the ROI mask), and then the ROI fluorescence had the background subtracted by a factor of 0.7. Each ROI was manually inspected to determine whether it was engram labelled or not using all z-stacks.

#### Analysis

All trials/sessions with motion in the *z*-axis were excluded from the analysis. The images were motion-corrected with suite2P. ROIs were also automatically detected with suite2P. Raw fluorescent data were exported into MATLAB (MathWorks) and analysed with custom-written scripts. Ca^2+^ responses were smoothed using a Savitzky–Golay filter with a 2^nd^ order polynomial and a 7-sample window and only transients longer than 250 ms were included in analysis. ROIs were first sorted into engram and non-engrams. Then, calcium events were detected when passing a threshold of 3 standard deviation from the median value of the baseline (measured with a rolling window over the pre -stimulus period) and peak amplitudes were measured using the ‘findpeak’ function in MATLAB. The values reported in the synchrony analysis represent the percentage of ROIs that were active at the same time within a window of 133 ms. A matrix containing binary vectors of calcium events for each ROI was built. For each trial, this matrix was summed along the ROIs dimension within a 133 ms window and then each value was divided by the number of ROIs. For control of the synchrony analysis, the vector containing the binarized calcium events was randomly shuffled.

### *Ex vivo* patch-clamp recordings

#### Slicing

Animals were deeply anaesthetized with isoflurane before being decapitated. The brain was quickly extracted (<1 minute) and placed in a carbogenated ice cold cutting solution containing (in mM): 110 Choline Chloride, 26 NaHCO3, 11.6 Na-Ascorbate, 7 MgCl2, 3.1 Na-pyruvate, 2.5 KCl, 1.25 NaH2PO4, 0.5 CaCl2 and 10 Glucose. Coronal slices (250-300 µm thick) of the auditory cortex were performed in ice-cold cutting solution with a vibratome (Leica Microsystems). Slices were left to incubate for up to 30 minutes at 35 °C, then at room temperature until recording in a carbogenated artificial cerebrospinal fluid (incubation aCSF) containing (in mM): 125 NaCl, 25 NaHCO3, 5 HEPES, 1 CaCl2, 6 MgCl2, 3 KCl, 1.25 NaH2PO4 and 10 Glucose.

#### Recordings

Whole-cell patch-clamp recordings were made from the soma of layer 5 pyramidal cells using an Olympus BX51 WI microscope equipped with a fluorescent imaging system (470 nm LED and 565 nm LED (Thorlabs), Retina 6000 camera (QImaging)) for the purpose of engram cells identification and dendritic visualization. Slices were continuously perfused with carbogenated aCSF at a rate of 2-3 mL/min at 34°C (± 1 °C). Borosilicate glass pipettes (inner diameter 0.86 mm, outer diameter 1.5 mm, Sutter) were pulled using an electrode puller (Sutter Instruments, USA) and had open tip resistances of 3–6 MOhm. Recording pipettes were filled with intracellular solution of the following composition (in mM): 130 K-Gluconate, 10 KCl, 10 HEPES, 4 Mg2+-ATP, 0.3 Na2-GTP, 10 Na2-Phosphocreatine (pH set to 7.25 with KOH, osmolarity 290 mosmol/L). 0.02% TMR biocytin (Thermo Fisher Scientific) was included in the intracellular solution for dendritic visualization and further cell identification with confocal imaging. Current clamp recordings were made using an amplifier (Multiclamp 700B; Molecular Devices, San Jose, USA) mounted on a remotely controlled micromanipulators (Luigs and Neumann, Germany). Cells were excluded from data analysis if the somatic resting membrane potential was more depolarised than -50 mV. Cells were also excluded if the somatic series resistance exceeded 50 MOhm or if the somatic series resistance changed by more than 10% during the recording (except when ZD7288 was applied). Membrane potential was not corrected for liquid junction potentials. Electrophysiological data were filtered at 10 kHz and acquired at 50 kHz by a Windows computer running Clampex 10.7 (Molecular Devices) using a Digidata 1440A interface (Molecular Devices, San Jose, USA). Data was analyzed using Clampfit 11.7

#### Optogenetic stimulations

To study the connectivity of engram cells, they were labelled with ChR2 (AAV_9_-TRE-EYFP-ChR2). As glial cells were also labelled with this construct and perturbed the visual identification of engram cells, slices were incubated in the incubation aCSF at 35°C with 0.2 µM of SR-101 (Sigma-Aldricht) for the first 15 minutes following slicing. This allowed to label astrocytes with a red fluorophore to avoid confusion with labelled neurons. Engram cells were confirmed to express functional ChR2 with a 500 ms 470 nm light pulse in current clamp and voltage clamp configurations. To evaluate synaptic inputs from engram cells onto the recorded neurons, light pulse (15 ms) was applied on the slice (10-15 repetitions). This protocol was performed before and after application of CNQX (10 µM, Sigma-Aldricht) and APV (100 µM, Sigma-Aldricht). 7/13 engram cells from this dataset were recorded simultaneously with a neighbouring non-engram cell.

#### Extracellular stimulation

Extracellular stimulation pipettes were pulled from theta glass capillaries (outer diameter 1.5 mm, Sutter) to approximate tip diameters of 1–5 µm. Both barrels were filled with recording aCSF connected to the poles of an isolated stimulator (WPI, Getting Instruments or ISO-Flex). The pipette tip was positioned under visual control in proximity (within 5 µm) of a tuft dendrite of the patched neuron (average 416.8 ± 22.69 µm from the soma, 193.8 ± 19.61 µm from the cortical surface). Brief pulses (0.1 ms) were used for different stimulation protocols (single pulse, trains of 2 pulses at 100 Hz of increasing intensities to evoke supralinear events, or trains of 5 pulses at 50 Hz (10-15 repetitions of the same intensity). The same protocol and intensities were performed before (control) and after ZD7288 (20 µM) application. All stimulations were performed under GABAa block (Gabazine 10 µm, Sigma-Aldricht).

### Statistical analysis

Unless otherwise stated, all statistical analysis was performed using Prism 10 software (Graphpad). For comparisons of two groups, t-tests were used to compare normally distributed data and Mann-Whitney or Wilcoxon tests were used to compare non-normally distributed data. For comparisons of more than two groups, one-way analysis of variance (ANOVA) with a Dunnett correction was performed. Values are reported as average +-SEM, and the statistical details of experiments are given in the captions of the figures. A threshold of p < 0.05 was used to define significance.

## Acknowledgements

We thank the Core Animal Services facility and the microscopy facility at the Florey Institute, Sharon Layfield for making the virus minipreps, and Michele Pignatelli for feedback on the manuscript. This work was supported by the Australian National Health and Medical Research Council (NHMRC) (APP2026307, L.M.P.; APP2011462, L.M.P. and T.R.; APP1130716, L.M.P. and T.R.) and Australian Research Council (ARC) (FT230100476, L.M.P.).

## Authors contribution

M.R. performed and analysed the behavioural, electrophysiological and histological experiments. L.G. and G.S. performed and analysed the imaging experiments. T.J.R. and L.M.P. supervised the project and advised on experimental design. M.R., T.J.R. and L.M.P. prepared the figures and wrote the manuscript.

## Supplemental Figures and Table

**Figure S1.**
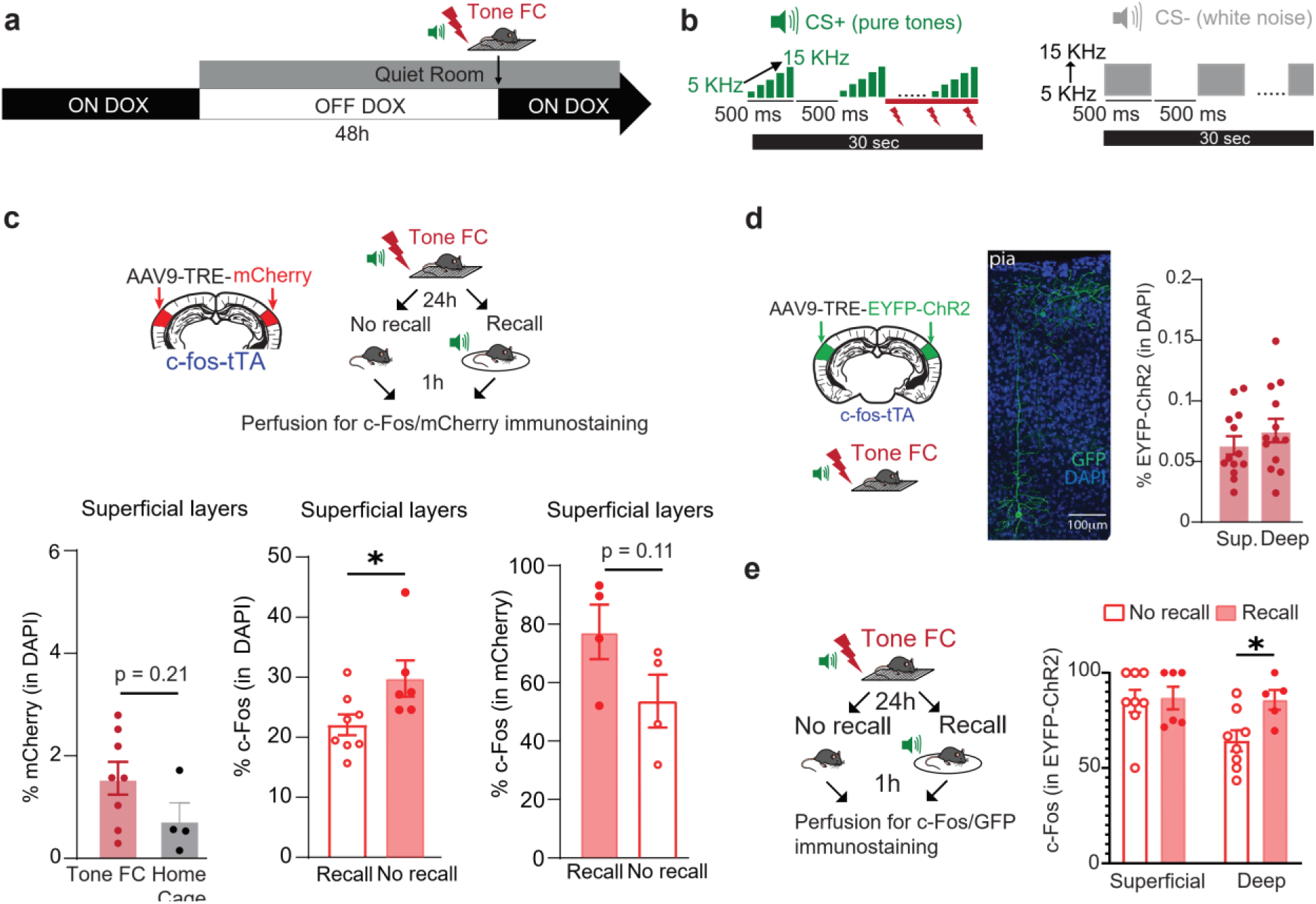
Engram cells labelling. **a,** Experimental timeline of engram labelling. **b,** Description of the CS+ and CS-. **c,** Engram cells labelling and recall-induced c-Fos expression in the superficial layers using the TRE-mCherry construct. A significant recall-induced c-Fos expression is observed in the superficial layers of the auditory cortex, but not in mCherry labelled neurons. No significant learning-induced labelling is observed in the superficial layers **d,** Engram cells labelling with the TRE-EYFP-ChR2 construct in the superficial and deep layers. **e,** Recall-induced c-Fos expression in EYFP-ChR2 labelled cells was observed only in the deep layers.

**Figure S2.**
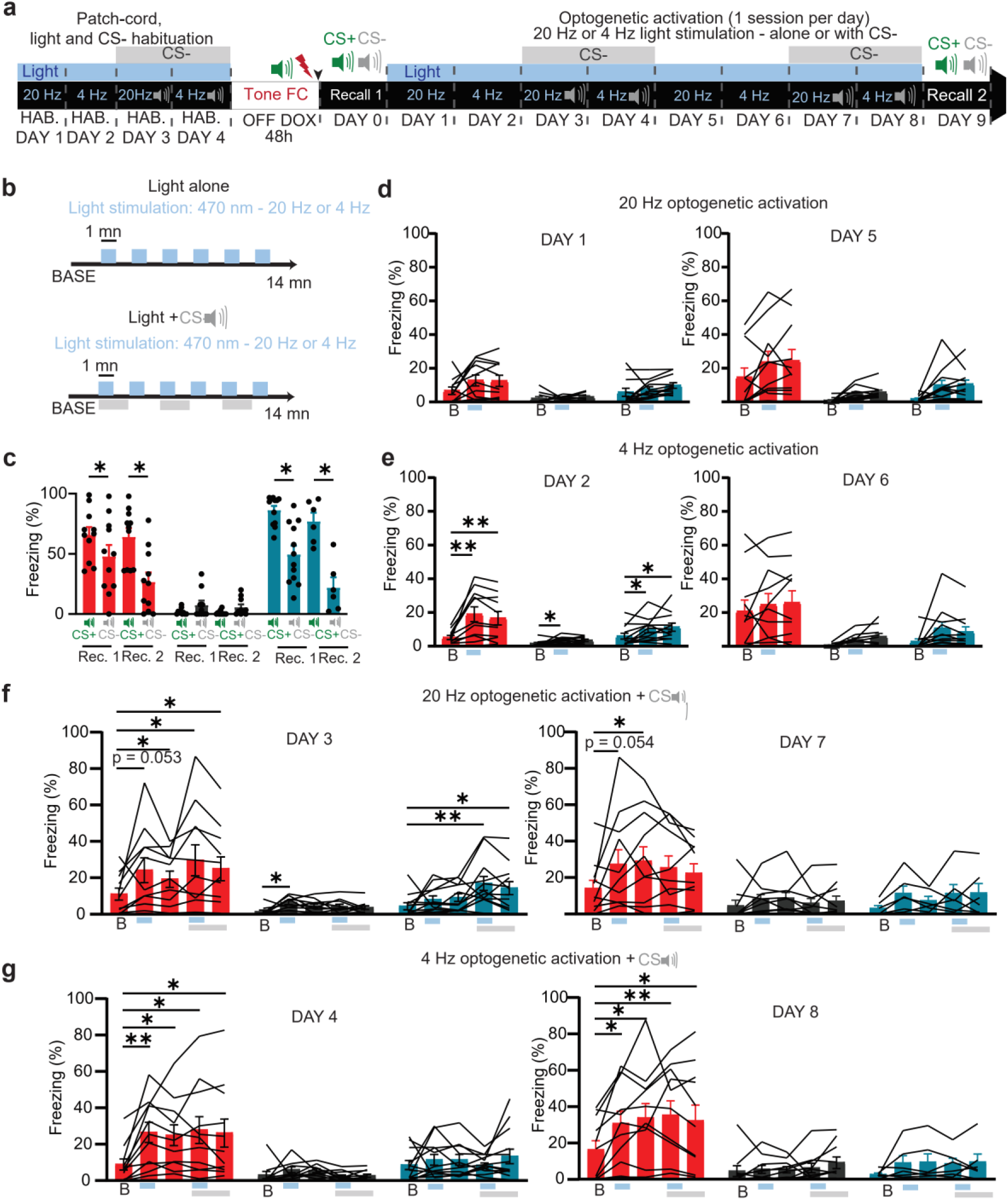
Optogenetic reactivation of engram cells. **a,** Experimental timeline for the gain of function experiment. Mice habituated to the patch-cord, light stimulation and the optogenetic context for 4 days before being put OFF DOX for engram cells labelling. Blue bars on top of the timeline represent sessions with blue light being delivered. Grey bars on top of the timeline represent sessions with the CS- being played. **b,** Protocol for optogenetic stimulation. Baseline lasted for 2 min, light ON and light OFF epochs lasted for 1mn. **c,** Freezing to CS+ and CS- during the first recall (DAY 0 on timeline) and the second recall (DAY 9 on timeline). **d-g,** Freezing quantifications for each of the 8 consecutive days of optogenetic reactivation of engram cells, grouped per condition (2 days per condition). Blue bars under the graphs represent light ON epochs, and grey bars the CS-. The 4 Hz + CS- condition is the only one evoking freezing in the experimental group only on the 2 days of optogenetic reactivation. Data are represented as mean ± SEM and significance was assessed with two-way ANOVAs with a Dunnett correction (*: p < 0.05, **: p < 0.01; ****: p < 0.0001).

**Figure S3.**
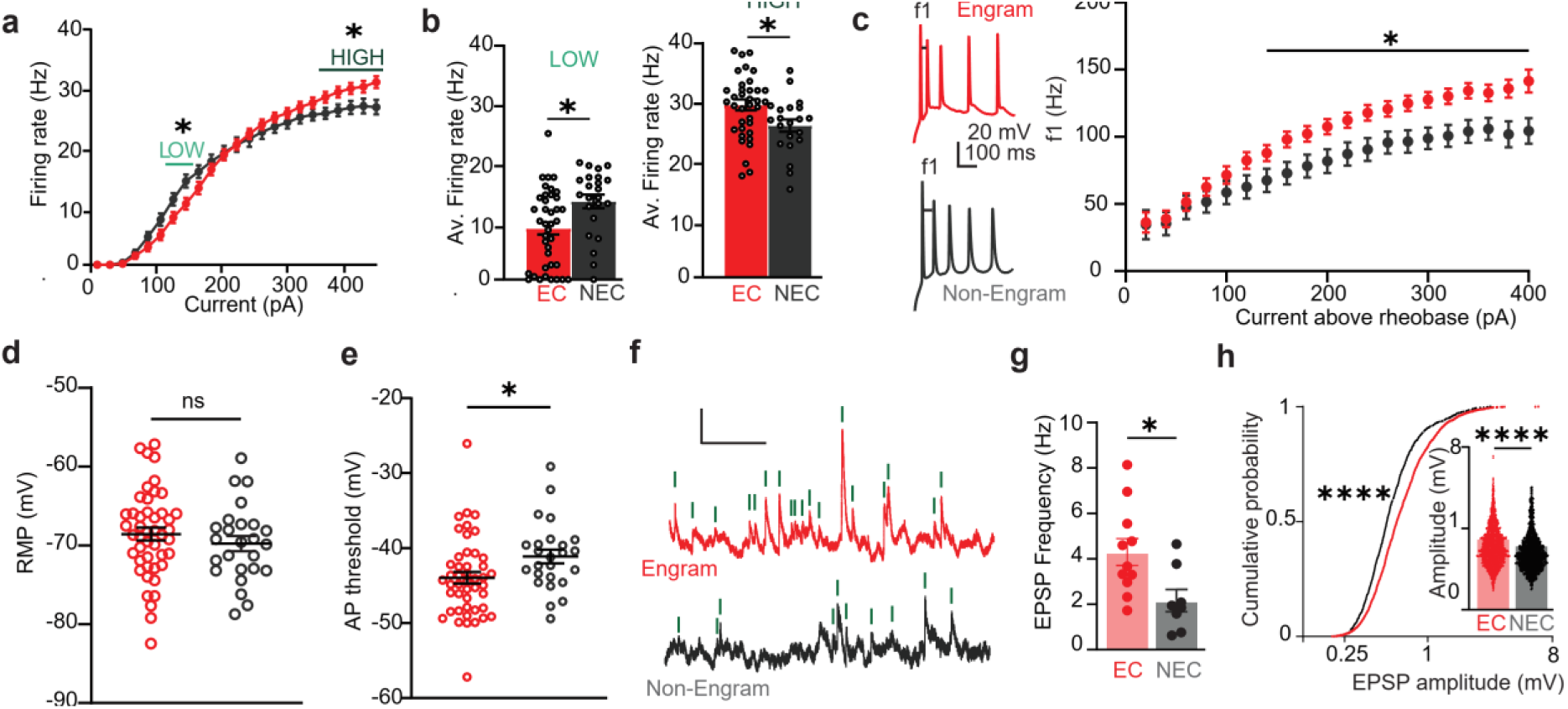
Intrinsic properties of EC 24h following fear conditioning. **a,** F-I curve showing the firing rate of EC and NEC in response to current step injections. A decreased firing rate is observed in EC for the 130-150 pA range of current injection (LOW), and an increase is observed for the 370-450 pA range of current injection (HIGH). **b,** Average firing rate over the low (left) and high (right) current injections. **c,** Initial frequency of firing (f1). Increased initial frequency in EC for high current injections (from 140 pA above rheobase) showing increased burst firing in EC. **d,** Resting membrane potential of EC and NEC. No difference was observed. **e,** Action potential threshold in EC and NEC. Significant decrease of AP threshold in EC. **f,** Example traces of spontaneous inputs on an EC labelled with mCherry (red, top) and a NEC (bottom, dark grey) recorded ex vivo 24h after conditioning (scale bars: 0.5 mV/1 sec). **g,** Average frequency of spontaneous EPSPs showing an increase in EC compared to NEC (EC: n = 2759 from 11 cells from 9 mice; NEC: 1253 events from 8 cells from 7 mice). **h,** Increased amplitude of spontaneous EPSPs in EC compared to NEC as shown per the cumulative distribution and the average. Data in the bar graphs are represented as mean ± SEM and significance was assessed with a two-tailed unpaired t-tests for a-e and h (*: p < 0.05; ****: p < 0.0001), two-tailed unpaired Mann-Whitney test for g (*: p < 0.05), and with a Kolmogorov-Smirnoff test for cumulative distribution (****: p < 0.0001).

**Figure S4.**
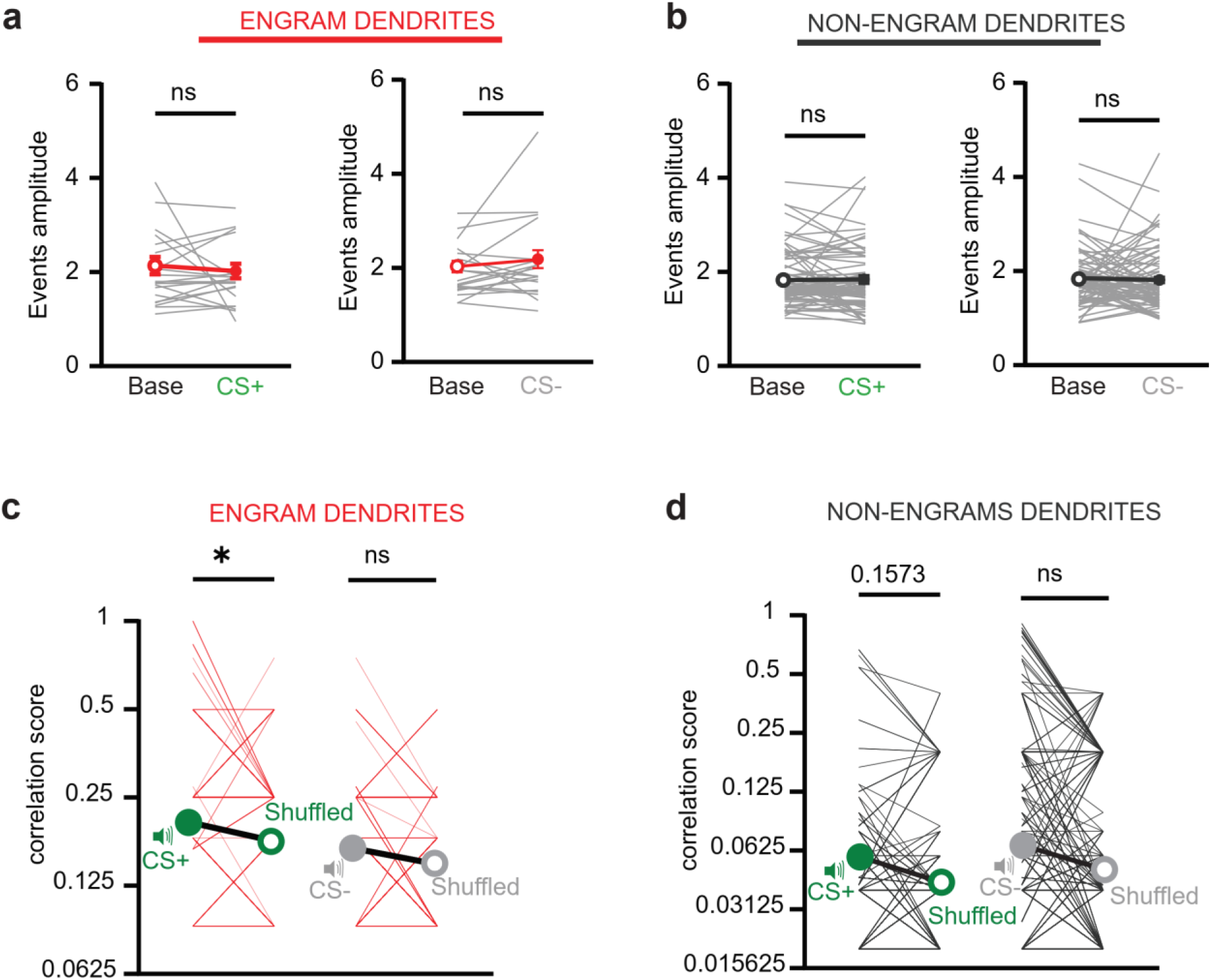
Two-photon imaging. **a-b,** Amplitude of dendritic calcium events in EC (a) and NEC (b) during CS+ and CS- exposures. No difference is observed. **c-d,** Comparison of the correlation scores with a shuffled dataset in EC (c) and NEC (d). Scales are logarithmic. A significant difference is observed between the EC dataset and the shuffled dataset for the CS+ data only. Data are represented as mean ± SEM and significance was assessed with two-tailed paired Wilcoxon tests (*: P < 0.05).

**Figure S5.**
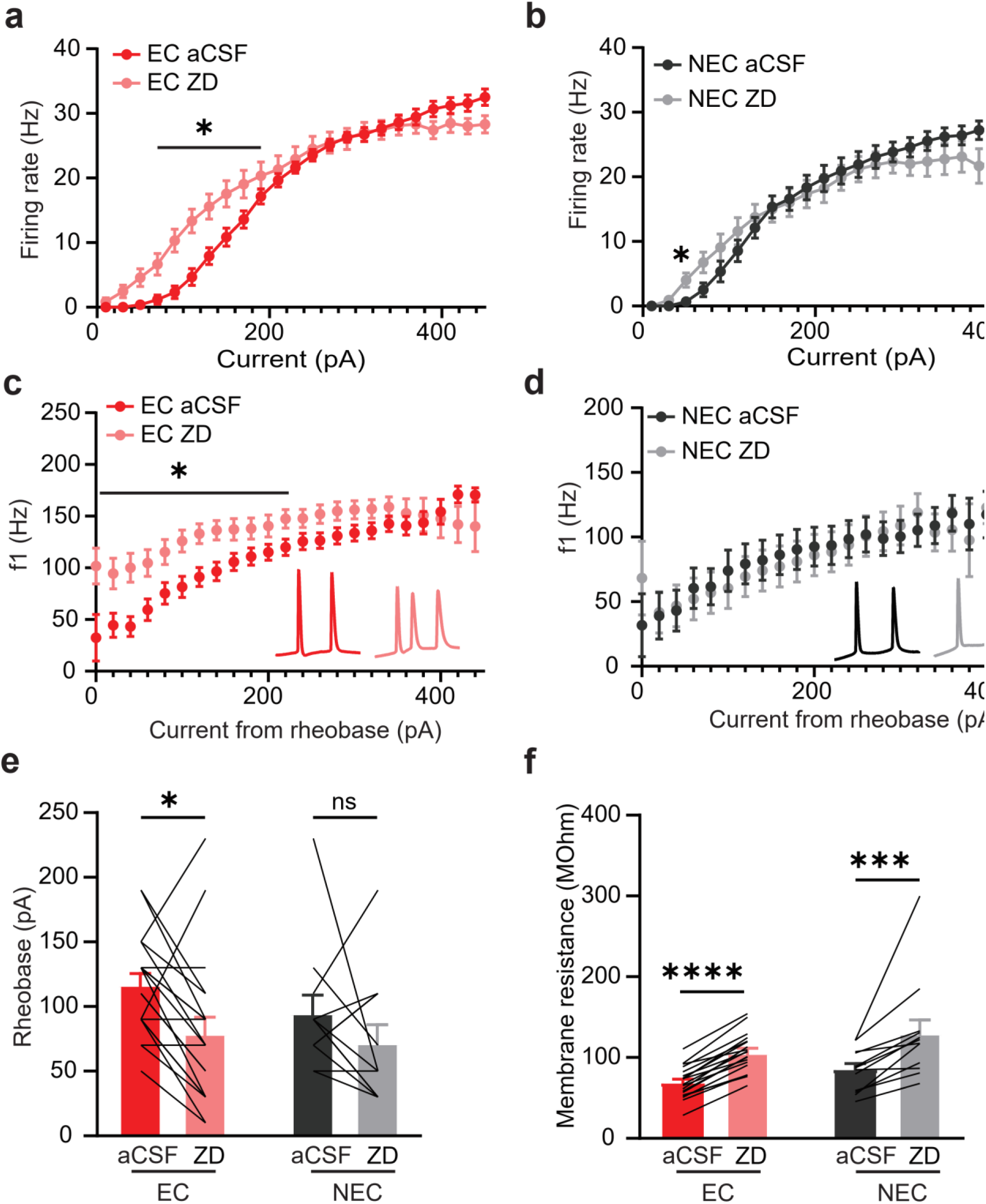
Increased effect of Ih block on EC excitability 24h following conditioning. **a-b,** F-I curves for EC (a) and NEC (b) showing the firing rate in response to current step injections without (aCSF) and with (ZD) Ih block. No effect of Ih block is observed at high current injections. **c-d,** Initial firing frequency (f1) in EC (C) and NEC (D) in response to current step injections without (aCSF) and with (ZD) Ih block. A significant increase of f1 under ZD is observed only in EC, from rheobase and until 220 pA above rheobase. **e,** Rheobase of EC and NEC without (aCSF) and with (ZD) Ih block. A significant decrease of the rheobase by Ih block is observed only in EC. **f,** Membrane resistance of EC and NEC without (aCSF) and with (ZD) Ih block. A significant increase of the rheobase by Ih block is observed only in both populations. Data are represented as mean ± SEM and significance was assessed with two-tailed paired Wilcoxon tests (*: p < 0.05; ***: p < 0.001; ****: p < 0.0001).

**Figure S6.**
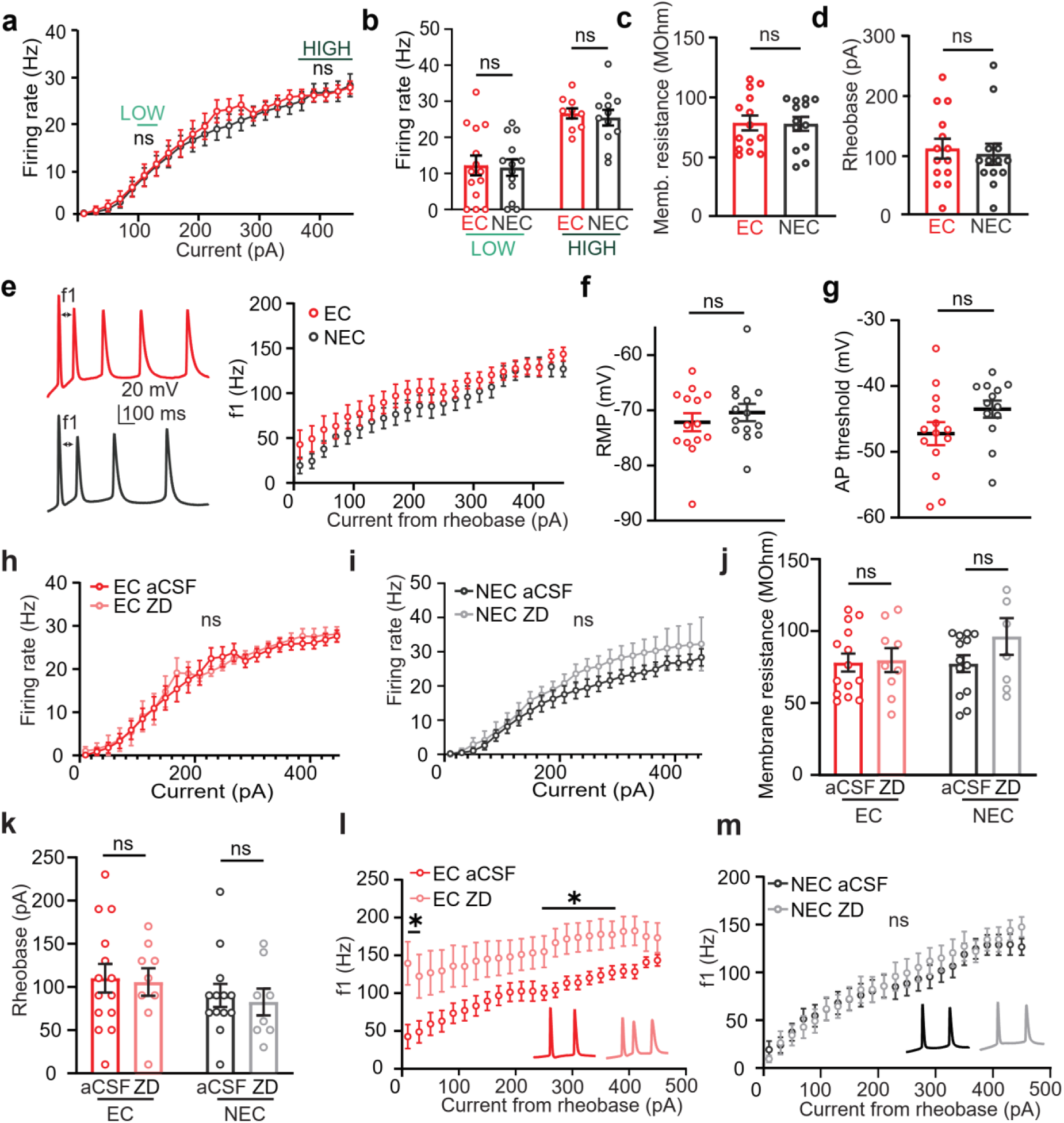
Intrinsic properties 1 week following conditioning and effect of Ih block. **a,** F-I curve in response to current step injections, showing no difference between EC and NEC at any current injection value. **b,** Average firing rate over the low (130-150 pA; left) and high (370-450 pA; right) current injection ranges, showing no difference between EC and NEC. **c,** Membrane resistance of EC and NEC showing no difference. **d,** Rheobase of EC and NEC showing no difference. **e,** Initial firing frequency (f1) of EC and NEC for each current step injection from rheobase and above. No difference was observed. **f,** Resting membrane potential of EC and NEC. No difference was observed. **g,** Action potential threshold in EC and NEC. No difference was observed. **h-i,** F-I curves for EC (H) and NEC (I) showing the firing rate in response to current step injections without (aCSF) and with (ZD) Ih block. No effect of Ih block is observed. **j,** Membrane resistance of EC and NEC without (aCSF) and with (ZD) Ih block. A significant increase of the rheobase by Ih block is observed only in both populations. **k,** Rheobase of EC and NEC without (aCSF) and with (ZD) Ih block. A significant decrease of the rheobase by Ih block is observed only in EC. **l-m,** Initial firing frequency (f1) in EC (l) and NEC (m) in response to current step injections without (aCSF) and with (ZD) Ih block. A significant increase of f1 under ZD is observed only in EC. Data are represented as mean ± SEM and significance was assessed with two-tailed unpaired Mann Whitney tests (*: p < 0.05).

**Table S1.** p-values corresponding to each session of optogenetic reactivation of engram cells. Each table summarizes the p-values for the comparison of the baseline freezing with each epoch during each session. In bold are shown the p-values for testing the average of the 2 sessions with the same experimental condition. Each table summarizes the p-values for each group. Significance is indicated with stars. Significance was assessed with two-way ANOVAs with Dunnett correction for multiple comparisons.

| SHOCK |  |  |  |  |  |
| --- | --- | --- | --- | --- | --- |
|  | Baseline vs | Light ON | Light OFF | Light ON+CS- | Light OFF+CS- |
| 20 Hz | Day 1 | 0.14 | 0.17 |  |  |
|  | Day 5 | 0.1 | 0.08 |  |  |
|  | **Average** | **0.07** | **0.06** |  |  |
| 4 Hz | Day 2 | 0.004* | 0.005* |  |  |
|  | Day 6 | 0.53 | 0.41 |  |  |
|  | **Average** | **0.077** | **0.1** |  |  |
| 20 Hz<br>🔊 | Day 3 | 0.053 | 0.04* | 0.035* | 0.024* |
|  | Day 7 | 0.054 | 0.03* | 0.01* | 0.22 |
|  | **Average** | **0.029\*** | **0.067** | **0.01\*** | **0.08** |
| 4 Hz<br>🔊 | Day 4 | 0.002* | 0.006* | 0.001* | 0.003* |
|  | Day 8 | 0.017* | 0.049* | 0.03* | 0.01* |
|  | **Average** | **0.018\*** | **0.012\*** | **0.004\*** | **0.014\*** |

| NO ChR2 |  |  |  |  |  |
| --- | --- | --- | --- | --- | --- |
|  | Baseline vs | Light ON | Light OFF | Light ON+CS- | Light OFF+CS- |
| 20 Hz | Day 1 | 0.37 | 0.08 |  |  |
|  | Day 5 | 0.12 | 0.1 |  |  |
|  | **Average** | **0.25** | **0.11** |  |  |
| 4 Hz | Day 2 | 0.024* | 0.049* |  |  |
|  | Day 6 | 0.21 | 0.37 |  |  |
|  | **Average** | **0.26** | **0.3** |  |  |
| 20 Hz<br>🔊 | Day 3 | 0.24 | 0.14 | 0.007* | 0.019* |
|  | Day 7 | 0.38 | 0.65 | 0.38 | 0.36 |
|  | **Average** | **0.75** | **0.81** | **0.053** | **0.15** |
| 4 Hz<br>🔊 | Day 4 | 0.55 | 0.7 | 0.85 | 0.25 |
|  | Day 8 | 0.49 | 0.51 | 0.48 | 0.55 |
|  | **Average** | **0.79** | **0.82** | **0.98** | **0.74** |

**Table S1. p-values corresponding to each session of optogenetic reactivation of engram cells.**
| NO SHOCK |  |  |  |  |  |
| --- | --- | --- | --- | --- | --- |
|  | Baseline vs | Light ON | Light OFF | Light ON+CS- | Light OFF+CS- |
| 20 Hz | Day 1 | 0.97 | 0.5 |  |  |
|  | Day 5 | 0.48 | 0.39 |  |  |
|  | **Average** | **0.8** | **0.64** |  |  |
| 4 Hz | Day 2 | 0.04* | 0.1 |  |  |
|  | Day 6 | 0.63 | 0.44 |  |  |
|  | **Average** | **0.78** | **0.66** |  |  |
| 20 Hz<br>🔊 | Day 3 | 0.013* | 0.078 | 0.065 | 0.13 |
|  | Day 7 | 0.63 | 0.57 | 0.86 | 0.74 |
|  | **Average** | **0.84** | **0.9** | **0.98** | **0.98** |
| 4 Hz<br>🔊 | Day 4 | 0.74 | 0.73 | 0.95 | 0.96 |
|  | Day 8 | 0.92 | 0.9 | 0.55 | 0.82 |
|  | **Average** | **0.99** | **0.99** | **0.99** | **0.99** |

